# GRIDSS, PURPLE, LINX: Unscrambling the tumor genome via integrated analysis of structural variation and copy number

**DOI:** 10.1101/781013

**Authors:** Daniel L. Cameron, Jonathan Baber, Charles Shale, Anthony T. Papenfuss, Jose Espejo Valle-Inclan, Nicolle Besselink, Edwin Cuppen, Peter Priestley

## Abstract

We have developed a novel, integrated and comprehensive purity, ploidy, structural variant and copy number somatic analysis toolkit for whole genome sequencing data of paired tumor/normal samples. We show that the combination of using GRIDSS for somatic structural variant calling and PURPLE for somatic copy number alteration calling allows highly sensitive, precise and consistent copy number and structural variant determination, as well as providing novel insights for short structural variants and regions of complex local topology. LINX, an interpretation tool, leverages the integrated structural variant and copy number calling to cluster individual structural variants into higher order events and chains them together to predict local derivative chromosome structure. LINX classifies and extensively annotates genomic rearrangements including simple and reciprocal breaks, LINE, viral and pseudogene insertions, and complex events such as chromothripsis. LINX also comprehensively calls genic fusions including chained fusions. Finally, our toolkit provides novel visualisation methods providing insight into complex genomic rearrangements.

## Introduction

Copy number alteration (CNA) and structural variation (SV) are two of the key classes of somatic mutations in cancer with approximately 90% of tumors undergoing significant rearrangement [1]. Despite the critical and interrelated role of SV and CNA in tumorigenesis [2], the mechanisms driving and consequences of genomic rearrangements in cancer are less well understood than for point mutation events, and less frequently used as biomarkers for clinical purposes. This is due both to the relative paucity of whole genome sequencing data which is required for SV analysis, but also the fact that genomic rearrangements have significant diversity and in many cases a high degree of complexity with individual events involving multiple or even up to hundreds of breaks [3, 4].

Notwithstanding the fundamental interconnectedness between CNA and SV breakpoint events, the vast majority of existing methods detect either SV breakpoints [5] or CNA [6] in isolation. Whilst SV callers incorporating read depth do exist [7], only a handful of methods perform a joint calling of SV and CNA [8, 9], none of which guarantee that the SV and CNA position match. Furthermore, existing SV tools typically don’t support all break junction types, requiring the use of specialised calling tools to find features such as viral insertions [10] and mobile line element insertions [11]. Critically, no existing caller utilises the single break-end reporting capability of the VCF specifications. This notation enables breakpoints to be represented where one side cannot be unambiguously placed and facilitates the partial resolution of variants in regions of the genome typically considered inaccessible to short read sequencing.

Downstream interpretation of structural variant breakpoints is also highly challenging and is important to both understanding mechanisms and reporting pathogenic impacts of genomic rearrangements. A number of methods have been proposed that cluster and chain structural variants to search for more complex event types such as chromoplexy or chromothripsis [3, 12] but none that do so in a comprehensive way across all variant types. Finally, whilst there are tools that can identify the underlying complex rearrangements responsible for gene fusions identified from RNA sequencing [13, 14], no tool exists that predicts gene fusions caused by complex genomic rearrangements from DNA sequencing alone.

Here, we present a comprehensive toolkit for integrated somatic copy number and structural variation analysis using short read whole genome sequencing data from tumor/normal sample pairs. Our toolkit consists of 3 highly integrated tools: GRIDSS version 2, an enhancement of the GRIDSS [15] SV caller that also performs SV chained assembly and reports single break-end variants; PURPLE, which performs SV-aware copy number segmentation and purity and ploidy estimation, and LINX, which clusters the genomic rearrangements into higher order events and interprets their impact.

We show that integrated SV and CNA analysis combined with accurate chaining, clustering and annotation of variants allows significant improvements in resolution of complex events, and highly sensitive and accurate prediction of clinically actionable events including gene fusion, homozygous deletion and amplification events.

## Toolkit Description & Results

### Somatic structural variation (GRIDSS)

The first component of the toolkit is version 2 of the GRIDSS structural variant caller [15]. Like the first version, GRIDSS performs genome-wide break-end assembly followed by probabilistic variant calling based on the totality of the direct read/read pair support, and the assembly contig support. As well as performance and output format improvements, GRIDSS version 2 adds somatic variant calling, structural variant phasing, and single breakend variant calling and classification capabilities.

Unlike the case of germline calling, no gold standard benchmarks exist for somatic variant calling and there is a lack of high quality truth sets for estimating precision and sensitivity of somatic SV callers. To fill this gap, we used a recently generated somatic SV truth set for the COLO829 tumor-normal cell line pair using a combination of Illumina, PacBio, Oxford Nanopore, and 10X Genomics sequencing, followed by targeted capture and PCR-based validations and manual curation (J.E.V., E.C., unpublished results). Based on benchmarking results [16, 17] we selected the widely used structural variant caller Manta [5] as a well-performing comparison tool. To test sensitivity and reproducibility, we ran both tools on 3 independent sequencing runs of the COLO829T/COLO829BL tumor/normal cell line pairs and used the truth set for determining false positive and negatives. On average, GRIDSS achieved a 93% sensitivity (88% for Manta), and it did so reporting 7 false positive calls (66 for Manta) (figure 2a, 2b). To validate the false positive rate, we repeated the experiment in triplicate with both ‘tumor’ and matched normal material sourced from the same cell line NA12878. For this negative control, GRIDSS reported on average only a single false positive event (45 for Manta) (figure 2c).

**FIGURE 1:**
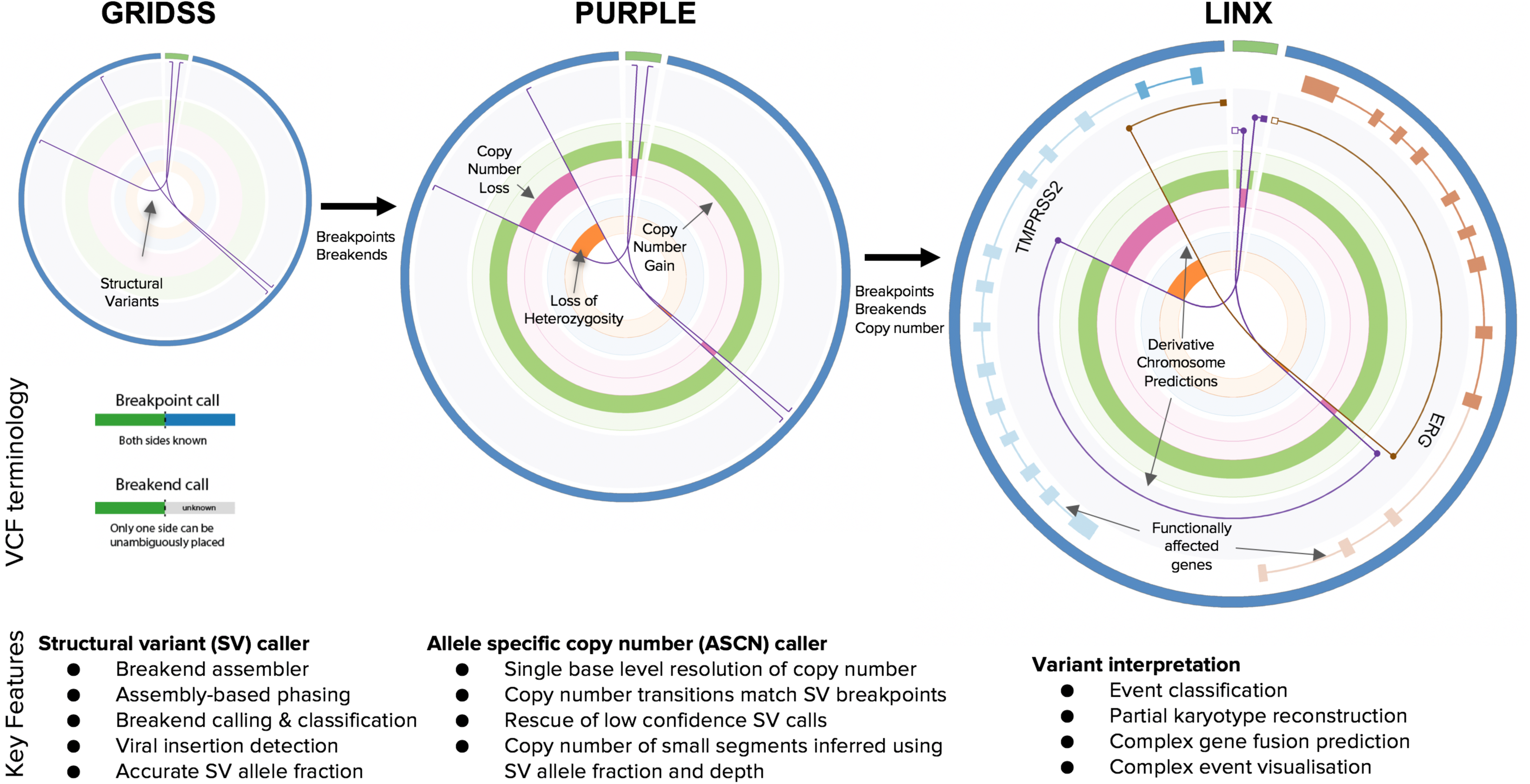
Toolkit Overview. Illustrative schematic of the 3 key components of the toolkit and the key features of each tool.

**FIGURE 2:**
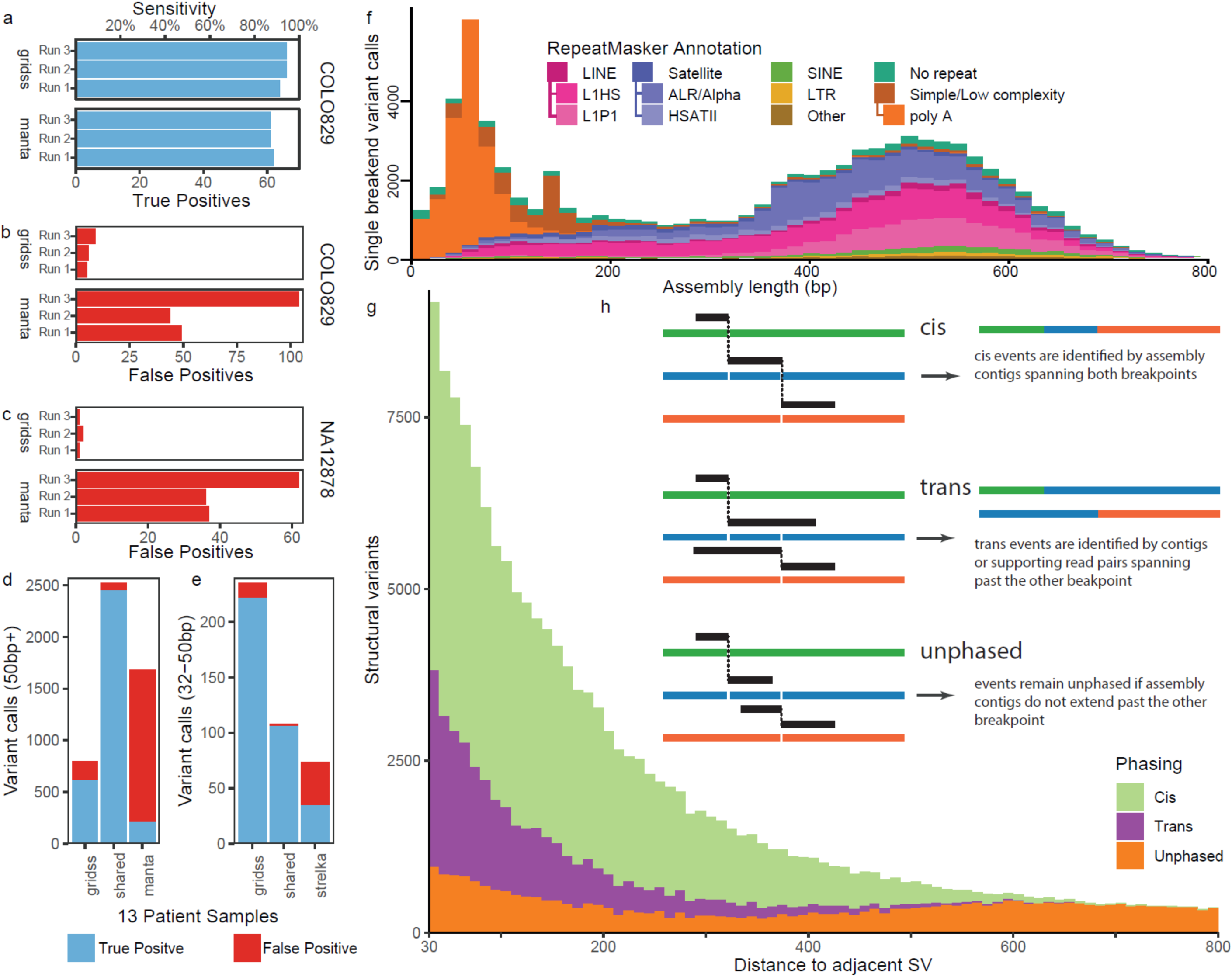
GRIDSS. a) Count of validated structural variant calls for GRIDSS and Manta on each of 3 biological replicates of the COLO829 paired tumor and normal cell lines. Sensitivity is displayed on the top axis. A structural variant truth set was created for COLO829 as part of the analysis using a multi-platform analysis including long read, short read, optical maps, and PCR analysis. b-c) Count of false positive variants for GRIDSS and Manta in each of 3 biological replicates for COLO829 (b) and a null tumor sample (c) respectively. The null tumor sample consists of independently sequenced NA12878 GIAB samples as both tumor and reference and hence has no somatic structural variants. d-e) Count of validated and not validated calls for structural variants across 13 patient tumor samples from the Hartwig Medical Cohort as validated by hybrid capture enrichment and sequencing. GRIDSS was compared to Manta for variants at least 50bp in length (d) and to Strelka for variants in the 32-50bp range (e). Shared refers to any variants that were found by both callers. f) Distribution of length and RepeatMasker annotation of assembled sequence of somatic single breakend variant calls across the Hartwig cohort. Only one side of single breakends can be unambiguously placed but the annotation of the assembled sequence is used classify the variant. The peak at 70bp is caused by poly-A truncated assemblies. The majority of somatic single breakends are LINE translocation or centromeric rearrangements. g) Phasing status of nearby somatic breakpoints in the Hartwig cohort. Phasing of structural variants from short read sequencing data is limited by the library fragment size length, and the minimum alignable sequence length (30bp for bwa-mem as used by GRIDSS). h) Structural variants are phased cis if a single assembly spans across both breakpoints, trans if the supporting assembles support the reference allele at the position of the other breakpoint, and remain unphased if the assemblies are too short.

We also performed independent validation of GRIDSS and Manta breakpoint calls from 13 patient tumor samples, using a hybrid capture approach with target probes flanking and overlapping break-junctions to independently validate over 5,000 calls identified by either or both tools. Again, GRIDSS outperformed Manta with 3,403 of 3,666 (93%) calls validated compared to 2,685 of 4,299 (65%) for Manta (figure 2d, supp. table 1). Of the private Manta calls not found by GRIDSS, just 230 of 1777 (13%) were validated compared to 836 of 1031 (81%) GRIDSS private calls. Whilst only precise GRIDSS calls passed filtering and were included in the call set, imprecise Manta calls had a similar validation rate (40/288, 14%) to Manta private calls with all validated imprecise manta calls called by GRIDSS precisely. Since the default minimum reported event sizes of GRIDSS and Manta are 32 and 50bp respectively, we compared 32-50bp events to the short indel caller, Strelka [18]. Again, in this range GRIDSS has superior calling performance with 329 of 343 (96%) calls validated against 142 of 182 (78%) Strelka calls (figure 2e). The 95% (219 of 232) validation rate of GRIDSS private 32-50bp calls compares favourably to the 47% (35 of 74) validation rate of Strelka private calls in this size range. Notably, GRIDSS finds many short DUPs of 32-100 bases which are largely missed by both Strelka and Manta (ext. figure 1b).

In addition to reporting breakpoints, GRIDSS has the unique ability to report *single breakend* variants. Single breakend variants are break junctions in which only one side can be unambiguously placed. For each single breakend variant, GRIDSS reports the assembled sequence thus providing insight into the sequence at the other side. This approach offers several advantages. Firstly, by aligning the assembled breakend sequences to a viral database, GRIDSS is able to detect retroviral integration sites and sequences, thus obviating the need to run specialised viral integration detection software. Secondly, it can classify single breakends by aligning back to the reference genome and annotating with the RepeatMasker repeat class. Running GRIDSS against the Hartwig Medical cohort of 3,524 matched tumor/normal WGS of metastatic patient samples [19], we find that somatic single breakends most frequently map to LINE repeat elements (30%), centromeric and peri-centromeric repeat sequences (18%) and low complexity sequences (42%), typically poly A (80% of low complexity sequences) (figure 2f). The presence of two single breakends in opposite orientation, one with a polyA sequence, the other with a LINE annotation enables us to classify LINE insertions in LINX. Similarly, a single breakend into an alpha satellite repeat is indicative of a break junction into centromeric sequence. Whilst in the general case we are unable to determine the correct candidate breakend partner location, in some cases, we were able to manually resolve the ambiguity to a copy number transition lacking a supporting SV. An example of this is the COLO829 single breakend into centromeric sequence that can only be matched to the centromeric copy number transition across chromosome 1. These capabilities represent a new state-of-the-art in the resolution of structural variants in repetitive regions commonly considered inaccessible to short read sequencing technologies.

The breakend assembly approach taken by GRIDSS also enables the assembly-based phasing of nearby variants. When two structural rearrangements occur in close proximity, a breakend assembly can span across both breakpoints (figure 2h) thus providing direct evidence that the two variants are adjacent on the derivative chromosome. Breakends closer than 30bp cannot be phased as BWA does not reliably align sequences shorter than this length. For the Hartwig Medical cohort, variants could be phased with GRIDSS up to around 500 bases, limited by the fragment size distribution of the sequencing library inserts. Fortunately, multiple nearby somatic structural variants are frequent with 24% of all structural variants in the 3,524 samples of the cohort having a variant between 30 and 1,000bp away. GRIDSS is able to phase 70% of these breakpoints with their closest structural variant (figure 2g). Of the phased variants, GRIDSS phases 72% of nearby variants on the same derivative chromosome (cis) and the remaining on separate chromatids (trans). This phasing information is critical to LINX as it exponentially reduces the number of possible paths through derivative chromosomes.

### Purity, ploidy & somatic copy number alteration (PURPLE)

The second component of the toolkit is PURPLE (PURity and PLoidy Estimator) which determines the allele specific copy number of every base of the genome.

Like many existing tools for whole genome copy number determination [6, 20], PURPLE divides the genome into segments of uniform read depth and b-allele frequency (BAF) and performs a grid search to determine the overall sample purity and ploidy which best fit the observed data. This is done in PURPLE via minimising a penalty function which weighs the relative plausibility of a tumor genome containing regions of subclonality against having significant deviation from a diploid genome. Some of the unique features in the PURPLE penalty function are additional penalties for solutions with major allele copy number less than 1 (since homozygous loss of large sections of the genome are unlikely) and explicit penalties for implausible situations in which the somatic SNV copy number exceeds the major allele copy number. Where a tumor is highly diploid and there is insufficient aneuploidy in the tumor sample to uniquely determine a purity, PURPLE will estimate the purity using somatic variant allele frequencies.

A key feature of the PURPLE algorithm is base pair consistency between the copy number and SV positions, allowing for base pair accurate copy number determination. Firstly, in the initial segmentation of copy number, uncertainty bounds are calculated for each copy number segment breakpoint, and segment boundary is adjusted to the SV position if one exists within the bounds. This includes single breakends called by GRIDSS, which allows accurate copy number breakpoint placement even if the SV cannot be fully resolved. Secondly, where multiple proximate SVs exist, PURPLE utilises the SV breakend variant allele frequency (VAF) and read count estimates provided by GRIDSS together with the copy number from flanking segments to infer the copy number of the short region. Thirdly, PURPLE also applies extensive copy number smoothing logic to remove residual GC bias and other noise, which may create false positive copy number segments. Finally, where there is no supporting breakpoint for a copy number segment, PURPLE will search a broader set of low confidence filtered GRIDSS SV calls and recover matching SV calls that may have been inadvertently filtered or lack the depth of support to be included in the initial high confidence call set.

Together, these measures give PURPLE a very high concordance between SV and copy number breakpoints. Across a large pan cancer cohort of 3,524 tumors, 96.2% of copy number transitions were supported by a structural variant breakend from GRIDSS with 0.6% of those rescued by PURPLE from low confidence calls. This base pair accurate concordance of copy number and structural variants has direct clinical relevance by allowing more accurate calling of homozygous disruption of tumor suppressor genes. Across the cohort we found that 4% of homozygous deletions of tumor suppressor gene drivers were caused by deletion smaller than 1kbp and a further 9% by a deletion of 1-10kbp, highlighting the importance of accurate and high-resolution copy number breakpoint determination.

PURPLE also makes significant efforts to accurately determine the allele specific copy number of every segment. PURPLE is packaged with a separate BAF tool, AMBER, which determines the BAF in the tumor sample of 1.3 million common germline SNP loci and uses those that are found to be very high confidence heterozygous in the germline sample. Once copy number segmentation is completed, the BAF observations for each segment in the tumor sample are adjusted for purity and used to estimate allele specific copy number. Unique to PURPLE, the allele specific copy number for each small copy number segment with no coverage is also inferred using the principle that only a single allele is likely to change copy number between adjacent proximate segments.

To evaluate performance and consistency of PURPLE, we compared it to ASCAT, and 2 other combined SV and CNV callers, CONSERTING and WEAVER, on the COLO829 cell line pair. Overall, the CN profile of ASCAT, PURPLE and CONSERTING showed strong concordance, with Weaver heavily over-segmenting on this dataset (figure 3a, supp. table 2).

**FIGURE 3:**
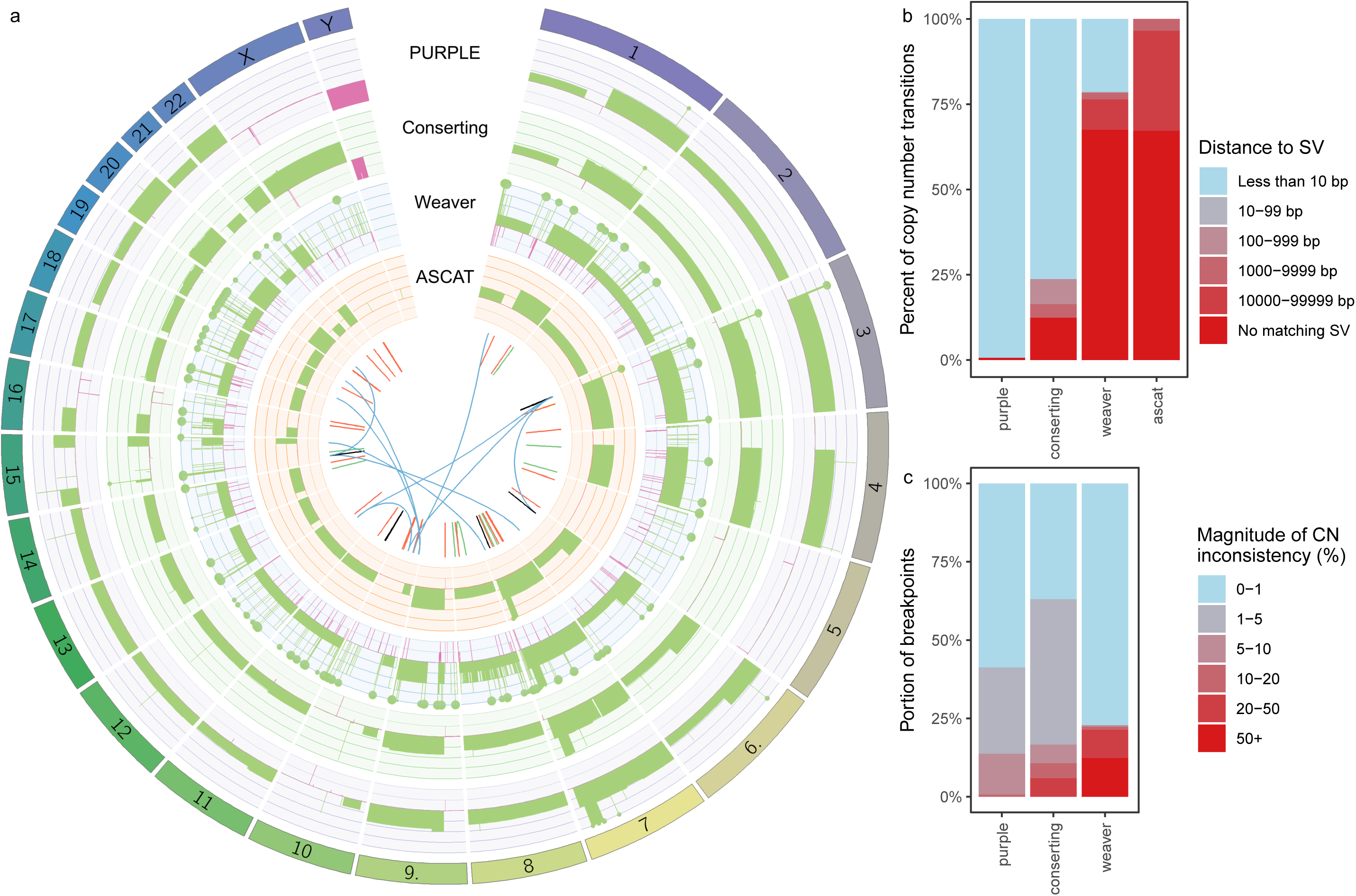
PURPLE. Benchmarking of copy number calling of PURPLE, CONSERTING, WEAVER, and ASCAT in the COLO829 cell line. a) A circos plot comparison of the copy number profile for each of the 4 callers. The scale for the copy number is 0 to 6 on each track with a filled circle at copy number 6 representing a segment with copy number of more than 6. Regions of copy number gain are shown in green and loss in magenta. The innermost circle shows the GRIDSS structural variant call set for COLO829T. b) Distribution of distances to nearest structural variant breakpoint across all copy number transitions for each caller. Copy number transitions occurring in centromeres or gaps in the reference genome were excluded. ASCAT was compared to GRIDSS calls. c) Distribution of difference in copy number change at either end of structural variant break junctions. A structural variant by definition causes the same copy number change at both ends so the difference should ideally be zero. Structural variants without corresponding copy number transitions on both sides were excluded.

We evaluated the internal consistency of copy number and SV breakpoints across the callers by comparing the position of the copy number and SV calls from PURPLE/GRIDSS, CONSERTING/CREST and WEAVER/WEAVER. Only PURPLE/GRIDSS generated an internally consistent SV/CNV call set with 98% of copy number transitions having a matching SV, compared to 75% for CONSERTING, and less than 25% for WEAVER (figure 3b).

For breakpoints having corresponding copy number transitions on both sides, the change in copy number on one side of the breakpoint should match the change in copy number on the other side. Using this consistency metric PURPLE/GRIDSS outperforms both CONSERTING and weaver (figure 3c), with the discrepancy between the copy number deltas on either side of breakpoints within 10% of the total copy number of the flanking segment for all break junctions. Whilst WEAVER has the highest portion of perfectly consistent copy number changes, this is due to the forcing of integer copy numbers and is offset by the high proportion of highly inconsistent transitions.

PURPLE/GRIDSS outputs directly correspond to a breakpoint graph with allele specific copy number and SV VAF annotations. This self-consistent representation is essential to sophisticated downstream analyses such as those performed by LINX.

### Clustering, Chaining, Annotation & Fusion Detection (LINX)

The role of the final component in the toolkit, LINX, is to group together the individual structural variant calls into distinct events, predict the local structure of the derivative chromosome and properly classify and annotate events to understand their mechanism and functional impact. There are several motivations for this, including reducing the complexity of interpretation of the overall genome rearrangement (in particular for shattering events and mobile LINE insertions), obtaining better insight into the frequency and characteristics of simple events, and better understanding of the nature and mechanisms in complex events. Gene fusion prediction sensitivity and accuracy can also be improved by comprehensive prediction of derivative chromosomes, especially in the case of complex rearrangements involving multiple clustered breaks.

LINX defines a genomic rearrangement ‘event’ as not just a single structural variant connecting 2 breakends, but as a group of 1 or more break-junctions which occurred proximately in time and transform the genome from one stable configuration to another. Events can be anything from a simple deletion or tandem duplication to a chromothripsis or breakage fusion bridge event. LINE insertions originating from a single source LINE element are classified together despite the fact that they may be continuing over the life of the tumor.

There are 4 key steps in the LINX algorithm. First, we annotate individual breakends with several basic geometric and genomic properties which are important to the clustering and chaining algorithm including whether each breakend is part of a foldback inversion, flanks a region of loss of heterozygosity, or is in a well known fragile site region [19]. LINX also uses a combination of previously known line element source information [21] and identification of both the local breakpoint structure and poly-A sequences to identify suspected mobile LINE source elements

Second, LINX performs a clustering routine. The fundamental principle for clustering in LINX is to join breakpoints where it is highly unlikely that they could have occurred independently. Rather than a single rule such as clustering variants into events based solely on proximity or variants which form a ‘deletion bridge’ [12], LINX relies on a set of 11 independent rules (see online methods). These include clustering variants which are very close in proximity (<5kb between breakends); clustering breakends which together bound an LOH or homozygous deletion; clustering translocations which share common arms at both ends; clustering inversion and long deletion & tandem duplication variants which directly overlap each other; and clustering all foldback inversions which occur on the same chromosome arm.

Third, after resolving all variants into clusters, LINX determines the derivative chromosome structures via a chaining algorithm. To do this, LINX considers all pairs of facing breakends on each chromosomal arm within each cluster and iteratively prioritises which pair is most likely to be joined. The chaining logic observes allele specific copy number constraints at all points on each chromosome and also the biological constraint that, unless strict criteria relating to detection of double minutes are met, circular chromosomes are not permitted without a centromere. Foldback and complex duplication replication events are explicitly modelled to allow chaining of clusters of variable ploidy and high amplification including double minutes (figure 4b) and breakage fusion bridge (ext. figure 2a). Overall, the chaining prioritisation scheme is designed to be error tolerant and aims to maximise the chance that each individual breakend is linked correctly to the next breakend on the derivative chromosome. However, due to multiple possible paths and upstream sources of errors, the prediction is representative only and unlikely to be completely correct in the case of highly complex clusters.

**FIGURE 4:**
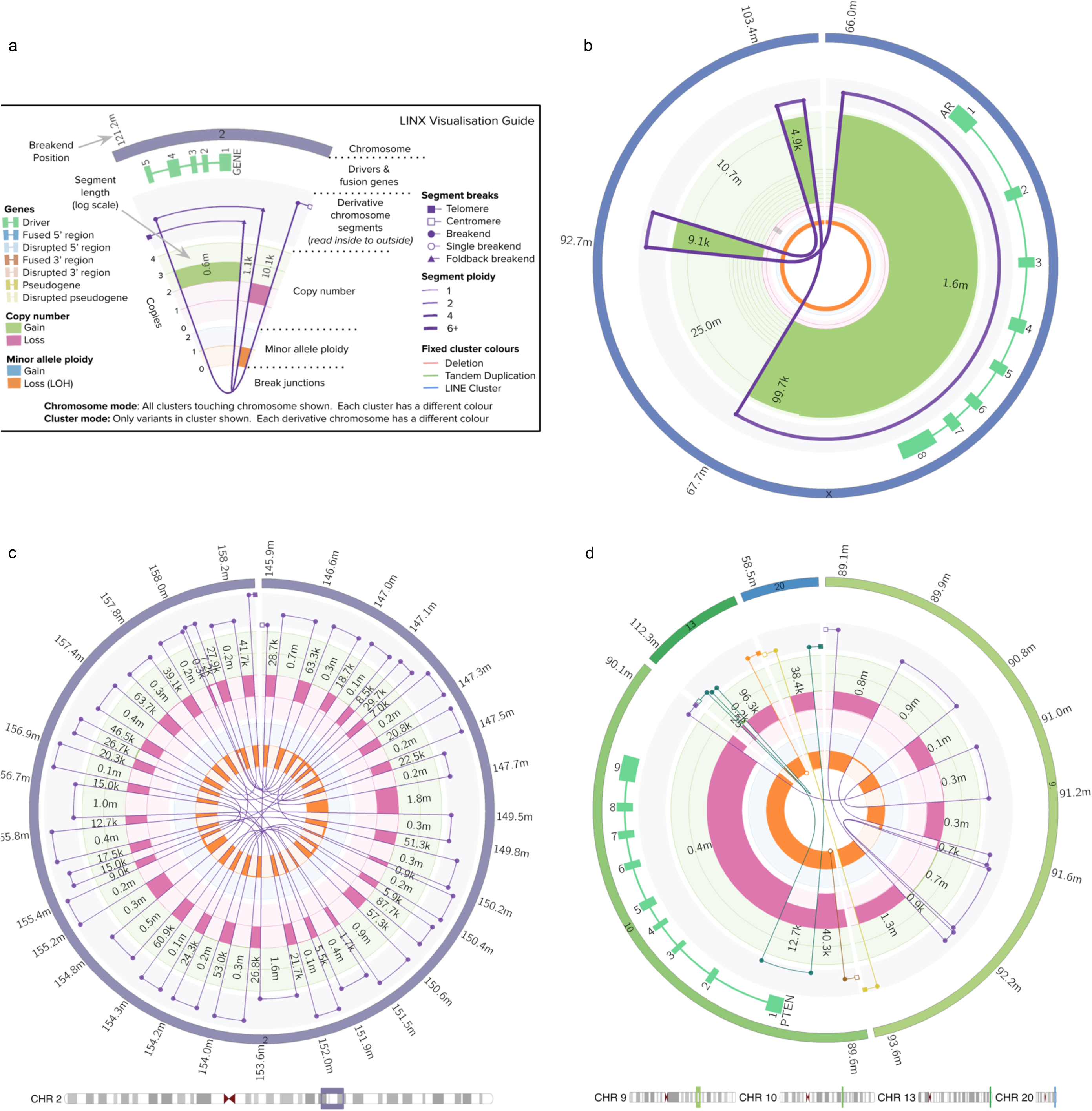
LINX. a) LINX Visualisation Guide b) Double minute formed from 3 variants in HMF004021A, a prostate tumor, and which amplifies known oncogene, AR, to a copy number of approximately 23x. c) Fully resolved chromothripsis event consisting of 31 structural variants affecting a 13mb region of chromosome 2 in HMF001571A, a prostate tumor. d) Chromoplexy like event in HMF001456A, a prostate tumor, consisting of 12 structural variants including 2 single breakends across 4 chromosomes. The event affects a region of broad loss of heterozygosity on chromosome 10 and causes a homozygous partial deletion of PTEN.

Once all clusters are fully chained, each cluster is also resolved with an annotated resolved type for all consistently chained one and two break junction events (ext. figure 3,4) with inconsistent one or two break junctions clusters marked as ‘INCOMPLETE’ (ext. figure 5) and clusters with 3 or more break junctions as COMPLEX. Resolution of event types in LINX can considerably simplify interpretation of the genome. In particular, isolating true simple deletions and duplications from variants which are notionally called as deletions and duplications but are part of a more complex event, allows clean length distributions to be discerned in simple events which are notably shorter than the distribution of deletion and duplication like variants in complex events. This improved distinction enables cleaner simple deletion and insertion profiles for downstream applications such as signature analyses and HR deficiency classification [22] which can be clinically relevant.

The fourth and final step in LINX is to annotate gene impact of breakends across all transcripts and predict gene disruptions and fusions. Gene fusions are called by searching for novel facing splice acceptor and donor pairs on the derivative chromosomes predicted by LINX with no limit on the number of breakpoints traversed. To meet the fusion calling criteria, the breakends must also connect to viable contexts in each gene and not be terminated by further breakends in the chain on either 5’ or 3’ partner end (see methods for full details).

To evaluate LINX fusion prediction, we compared LINX fusions to STAR-Fusion [23] predictions across 534 samples with matching whole genome and whole transcriptome data. Overall, LINX predicted 84% of 1,478 in-frame fusions detected by STAR-Fusion (supp. table 3). Whilst 86% of LINX intergenic fusion predictions are not detected in RNA expression, this is unsurprising as many rearrangements may be passenger events (like most point mutations) and, as the average expression of 50% of genes is less than 20 transcripts per million [24], fusions in genes with low of no expression are undetectable by RNA-seq.

For clinically relevant pathogenic and likely pathogenic fusions, LINX is highly sensitive. On the same 534 samples, amongst 393 known pathogenic fusion gene pairs [25-27] LINX predicted 36 out of 37 fusions from STAR-Fusion (figure 5a,s supp. table 3) and a further 11 putative fusions not called by STAR-Fusion, 5 of which have at least 1 supporting RNA-Seq read. LINX also predicted 24 of 26 fusions detected by STAR-Fusion involving genes from a curated list of 67 promiscuous 3’ and 5’ clinically relevant fusion gene partners, and also 50 additional promiscuous fusion candidates not detected in RNA. Altogether, 21 of the 130 (16%) known and promiscuous fusion predictions in the 534 samples were chained fusions (5 of which were confirmed in the RNA-seq data), highlighting the utility of chaining of derivative chromosomes.

**FIGURE 5:**
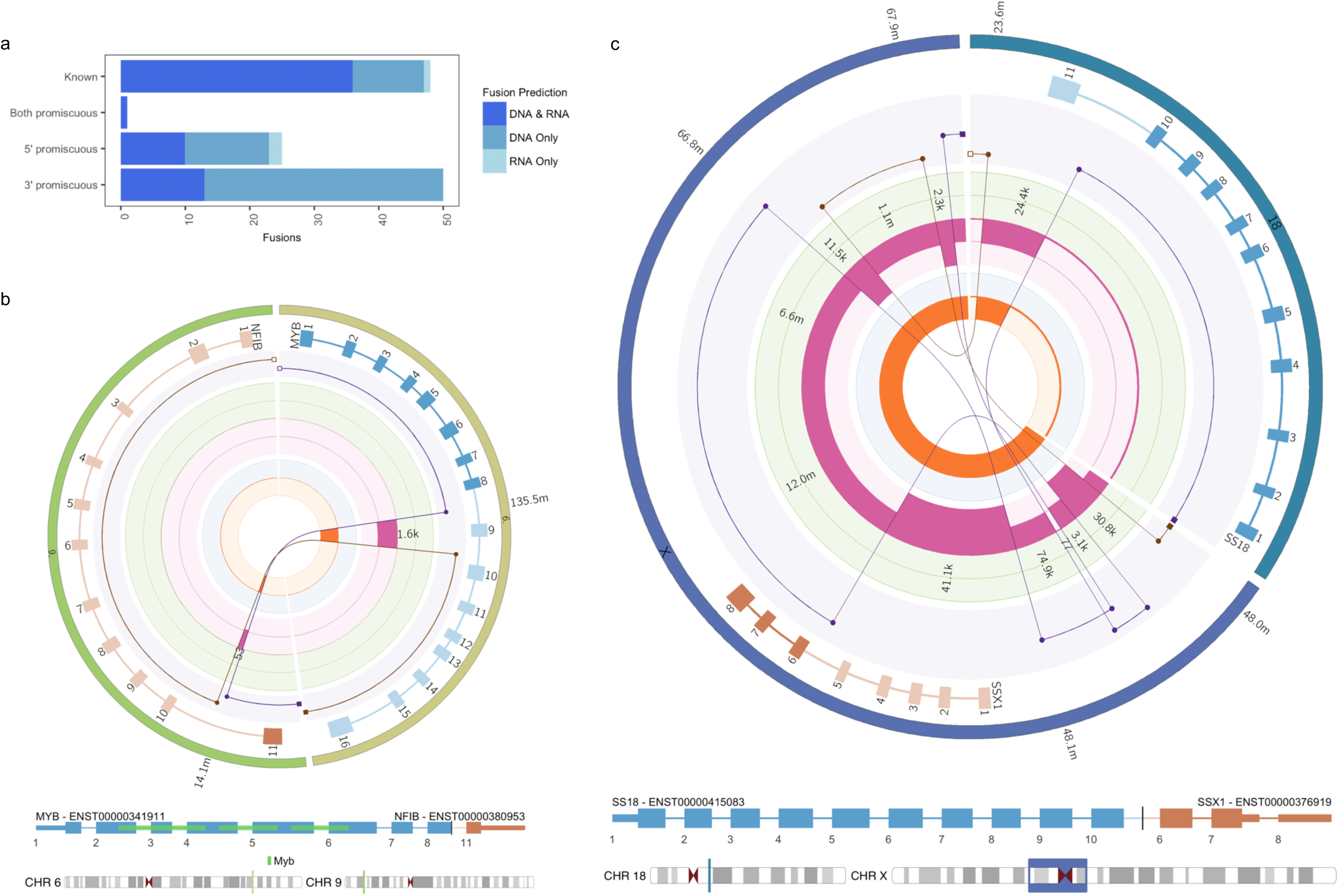
Fusions in LINX. a) Comparison of LINX fusion predictions to STAR-Fusion calls for pathogenic or potentially pathogenic fusion pairs in 534 tumor samples with matching whole genome and whole transcriptome data. Fusion predictions are classified as known if previously identified as pathogenic or as promiscuous on either the 5’, 3’ or both sides for fusions involving a gene that has previously been identified in at least 3 other fusions combinations. b) A MYB-NFIB fusion caused by a reciprocal translocation in HMF000780A, a salivary gland tumor. The translocation links exons 1-8 in MYB to exon 11 in NFIB. c) A SS18-SSX1 chained fusion caused by a complex cluster of 6 variants in HMF003579A, a Sarcoma. The predicted derivative chromosome links exons 1-10 in SS18 to exons 6-8 in SSX1 via a chain of 2 structural variants.

Finally, LINX provides novel detailed visualisations of the rearrangements in the tumor genome (fig 4a) either by cluster or chromosome, creating an integrated Circos [28] output showing copy number changes, clustered SVs, the derivative chromosome predictions and impacted genes including protein domain annotation for gene fusions all on the same diagram (figure 5b,5c). The visualisations use a log-based position scaling between events so that small and large scale structure can both be inspected on a single chart, allowing unprecedented resolution of complex structures across a broad array of events ranging from relatively simple reciprocal translocations (ext. figure 2c) and deletions with shards to complex structures such as chromothripsis (figure 4c), chromoplexy (figure 4d), multiple LINE insertions from the same source element (ext. figure 2c), pseudogene insertions (ext figure 2d), and other complex events which may even cause multiple drivers simultaneously (ext. figure 6).

## Discussion

Cancer genomes are often highly rearranged due to a mix of processes that impact intermittently (e.g. chromothripsis, BFB) or continuously (e.g. DNA repair deficiency) on the integrity of the tumor cells genome. Due to the complexity of dissection of all such events through whole genome sequencing, there are many potential sources of error which can confound correct interpretation of these genomic rearrangements including sample preparation, sequencing errors and biases (such as GC bias), inaccurate fitting of sample purity and ploidy, false positive or false negative structural variant calls, and inaccurate local copy number measurement. Accurate interpretation of structural variation is highly sensitive to all inputs, and error tolerant algorithms are essential to ensure that inaccuracies do not compound on each other.

GRIDSS, PURPLE and LINX together provide an integrated toolkit of algorithms that combines highly accurate structural variation and copy number calling with novel interpretation and visualisation methods to give unprecedented resolution into tumor genomic rearrangements using standard short read sequencing as input. Each of these tools possess novel features that make critical contributions to the final results. Particularly important are single breakend variant calling, phasing of nearby structural variants, classification of single breakend sequences, base-pair positional consistency between structural variant calls and copy number segmentation, rescue of low confidence structural variants, inference of copy number for short segments from structural variant depth and allele fraction, complex fusion prediction, and event classification/partial karyotype reconstruction.

The breakend assembly approach taken by GRIDSS enables assembly-based phasing of structural variants. With 11% of all somatic structural variants able to be phased, this represents a considerable reduction in the exponential number of possible paths and makes a significant contribution to LINX’s derivative chromosome prediction. Similarly, single breakend variant calls will typically be supported by a breakend assembly hundreds of base pairs long. These sequences are long enough that integrated viruses can be identified, and a reliable classification of the repeat type can be made. The ability to accurately identify and classify these structural variants, gives insights into regions of the genome typically considered inaccessible to short read sequencing technologies.

Whilst the toolkit has been optimised for short read technology, the short read length is ultimately the key limitation since it limits the phasing of proximate variants. In practice, LINX is able to resolve many structures via various chaining and clustering heuristics, but for more complex events, particularly highly rearranged focal regions, errors are inevitable and the chaining is only partial and representative. Derivative chromosome prediction can be improved with long read sequencing technologies. For example, 20kbp reads would allow an additional 6% of structural variants in the Hartwig cohort to be phased and allow a greater portion of LINE translocations to be fully resolved. Unfortunately, long reads are unable to resolve the telomeric and centromeric breakend ambiguities and still require supplementary approaches such as optical mapping [29] to fully resolve derivative chromosomes. At this time, cost and the high indel error rate of long reads makes short read technology an appropriate choice for comprehensive somatic variant determination for routine patient diagnostics and stratification towards precision medicine as well as for biomarker discovery

Despite the limitations of short reads, the clinical utility of the combined toolkit is significant. Detection of viral integrations, such as HPV, can directly impact treatment [29]. Base level accurate allele specific copy number in PURPLE allows more sensitive calling of regions with LOH and homozygous deletions and allow a more accurate classification of bi-allelic inactivation driven by short copy number segments. Indeed, PURPLE, if provided with somatic SNVs, will predict both the biallelic status and subclonal likelihood of each point mutation, aiding in the interpretation of variants of uncertain significance. For fusion calling, LINX provides comprehensive annotation of both simple and chained fusions and can achieve similar sensitivity to recognised RNA-seq methods for clinically relevant fusions.

Whilst the quality of the sequence data is often overlooked, it is a critical part of the overall pipeline. GRIDSS/PURPLE/LINX have been optimized for Illumina HiSeqX and NovaSeq sequencing data using fresh frozen samples, 100/40x tumor/normal coverage, and a 500-600bp library fragment size distribution. High quality sequencing data is critical to the high sensitivity and low FDR achieved by this toolkit.

This toolkit lays the foundation for answering numerous open questions about cancer evolution, DNA damage/repair, and somatic rearrangements more generally.

## Methods

See accompanying online methods.

## Data availability

All raw (BAM), analysed (VCF, SV, purity copy number data) germline and somatic genomic data used in this study was obtained from the Hartwig Medical Foundation (Data request DR-005). Standardized procedures and request forms for access to this data can be found at https://www.hartwigmedicalfoundation.nl/en.

Raw and analyzed data for the creation of the COLO829T/COLO829BL tumor/normal cell line pair structural variant truth set are available grouped under ENA study accession PRJEB27698. COLO829T copy number validation and capture panel validations of 13 tumor samples have been submitted to EGA (accession numbers pending).

## Code availability

GRIDSS, PURPLE and LINX are freely available as open source software from the Hartwig Medical Foundation (https://github.com/hartwigmedical/) and the Walter Eliza Hall Institute of Medical Research (https://github.com/PapenfussLab/gridss) under a GPLv3 license. A combined docker image is available from https://cloud.docker.com/u/gridss/repository/docker/gridss/gridss-purple-linx. Reference data required to run GRIDSS/PURPLE/LINX toolkit on hg19 is available from https://resources.hartwigmedicalfoundation.nl.

## Acknowledgements

This publication and the underlying study have been made possible partly on the basis of the data that Hartwig Medical Foundation and the Center of Personalised Cancer Treatment (CPCT) have made available to the study.

## Extended Figure Legends

**EXTENDED FIGURE 1:**
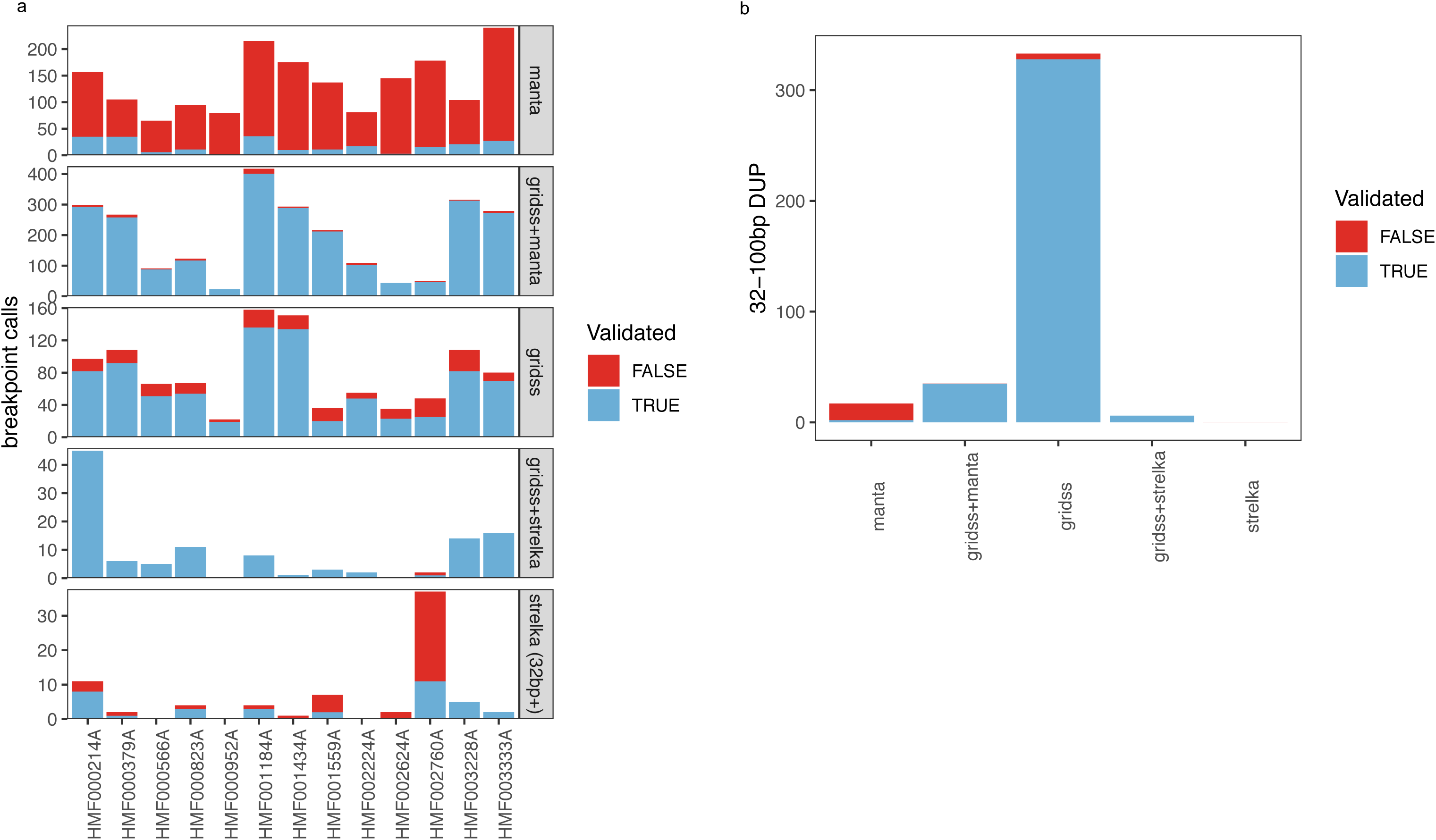
a) Count of validated and not validated calls for all structural variants for each of 13 patient tumor samples from the Hartwig Cohort as validated by capture panel. GRIDSS and Manta were compared for any variants > 50 bases in length. For short deletion and tandem duplications of >32 bases and <=50 bases GRIDSS was compared to Strelka. b) Count of validated and not validated calls for all tandem duplications of 32-100 bases in length across 13 patient tumor samples.

**EXTENDED FIGURE 2:**
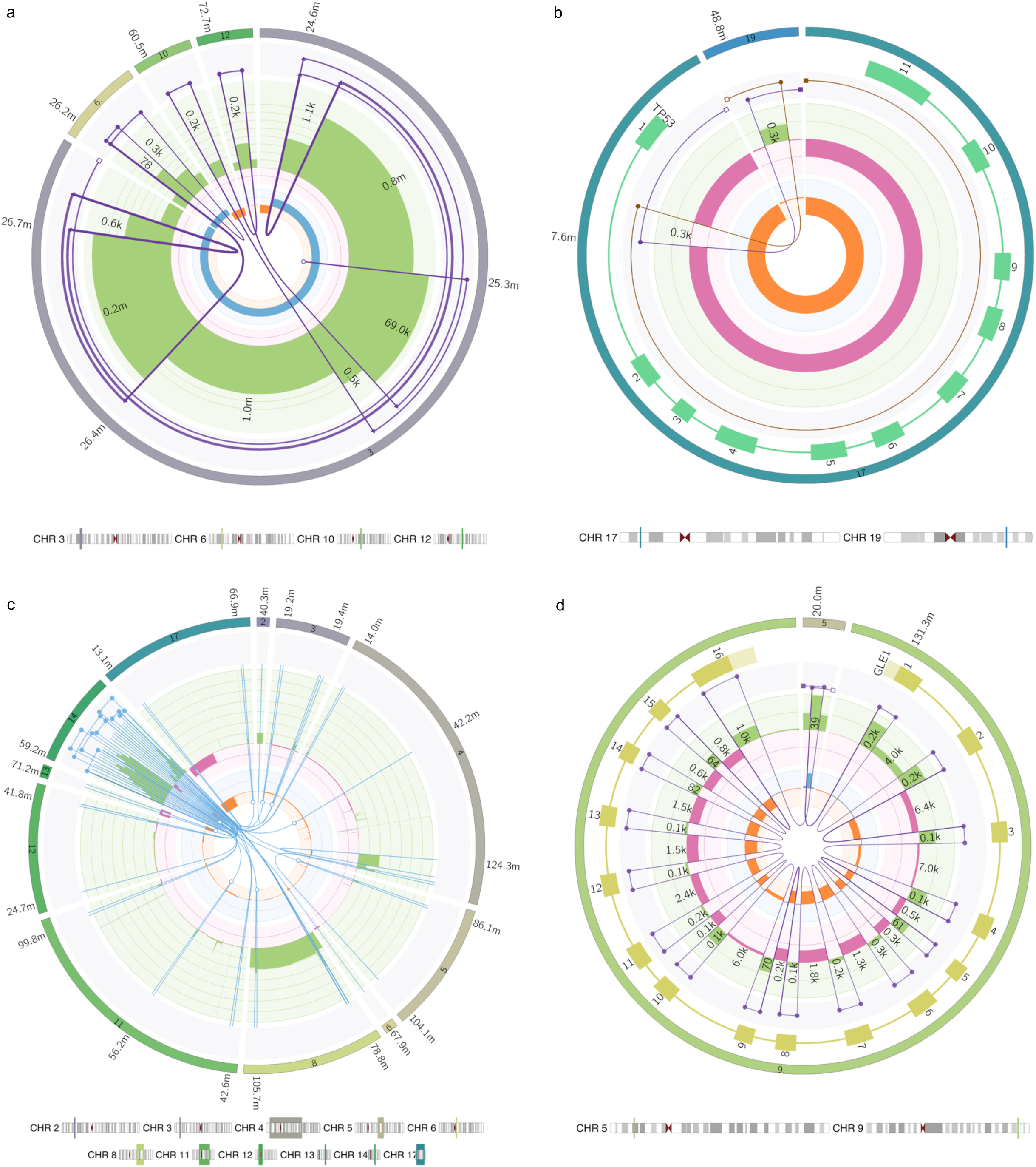
a) Breakage fusion bridge event affecting the P arm of chromosome 3 in the cell line COLO829T. The predicted derivative chromosome has a copy number of 2 and can be traced outwards starting from the centromere on chromosome 3, traversing 2 simple foldbacks and 2 chained foldbacks and finishing on a single breakend at chr3:25.3M which from the insert sequence can be inferred to be connected a centromeric satellite region (likely chromosome 1 which has a copy number gain of 2 over the centromere from P to Q arm, and which appears to be connected to chromosome 3 in unpublished SKY karyotype figures http://www.pawefish.path.cam.ac.uk/OtherCellLineDescriptions/COLO829.html). One chained foldback at chr3:26.4M includes a genomic shard from chr 6 of approximately 400 bases which has itself been replicated and internally disrupted by the foldback event. The other chained foldback at chr3:25.4M includes 2 consecutive genomic shards inserted from chromosome 10 and 12 of approximately 200 bases each. b) Reciprocal translocation affecting TP53 in HMF001913A, a prostate tumor. The 2 predicted derivative chromosomes overlap by approximately 300 bases on both ends, but are trans phased which rules out the possibility of a templated insertion at either location. Although the TP53 copy number alternates between 1 and 2, no derivative chromosome contains the full gene and the gene is homozygously disrupted. c) Complex LINE cluster in HMF002232B, a colorectal cancer. Overlapping segments from the LINE source element from chr 14:59.2M has been inserted at least 20 times independently at locations scattered throughout the genome. d) Pseudogene insertion of GLE1 into an overlapping break junction on chromosome 5 in HMF002165A, a non-small cell lung cancer. All 16 exons of the GLE1 canonical transcript are inserted, but part of the first and last exons are lost.

**EXTENDED FIGURE 3:**
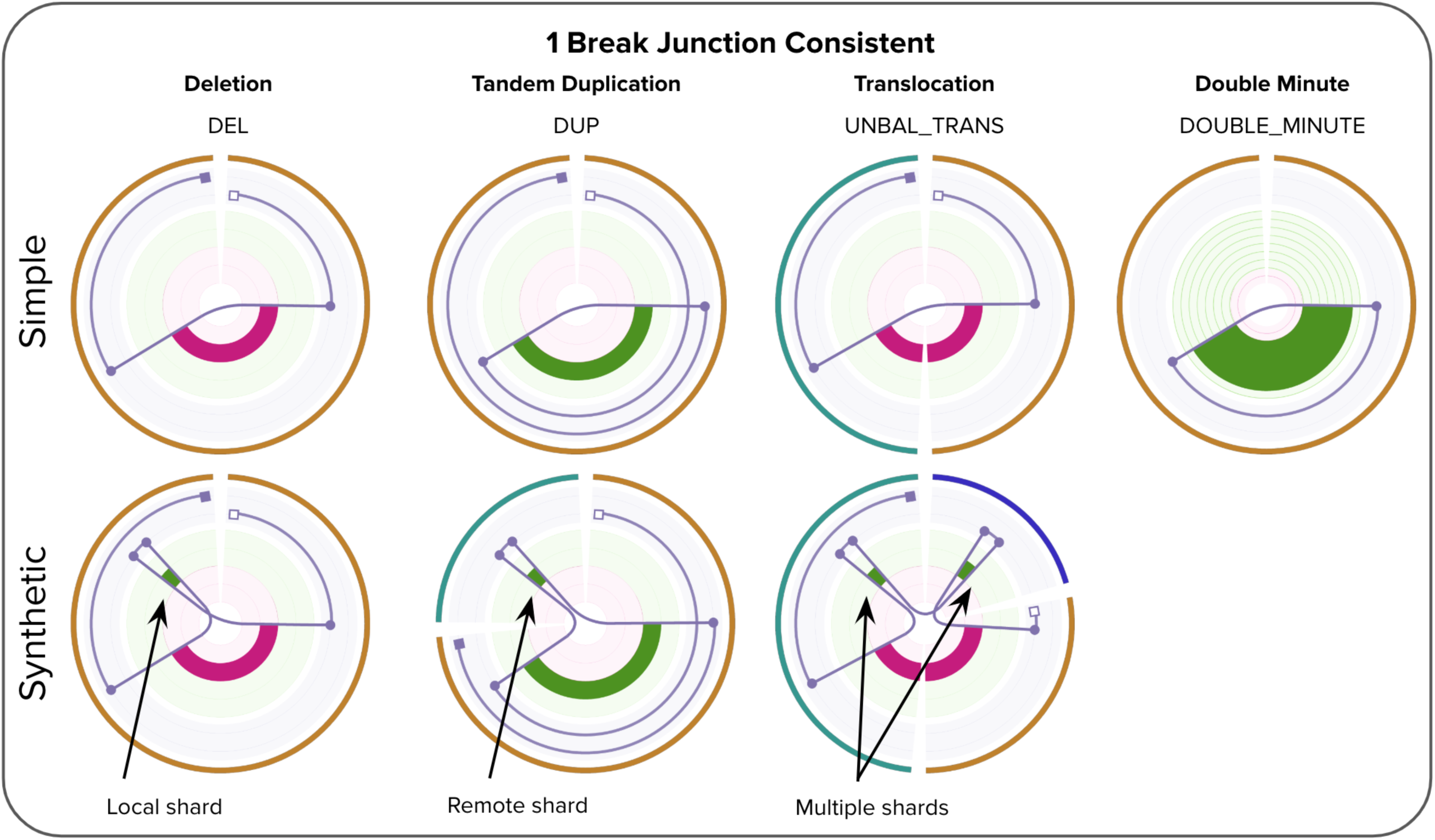
Chromoplexy like cluster formed from 19 break junctions across 7 chromosomes in HMF001596B, a prostate tumor. The rearrangement leads 3 distinct putative drivers in a single event, including a chained TMPRSS2-ERG fusion with 2 hops, a loss of heterozygosity for PPP2R2A which also has a stop gained point mutation (not shown), and an intronic homozygous disruption of PTEN.

**EXTENDED FIGURE 4:**
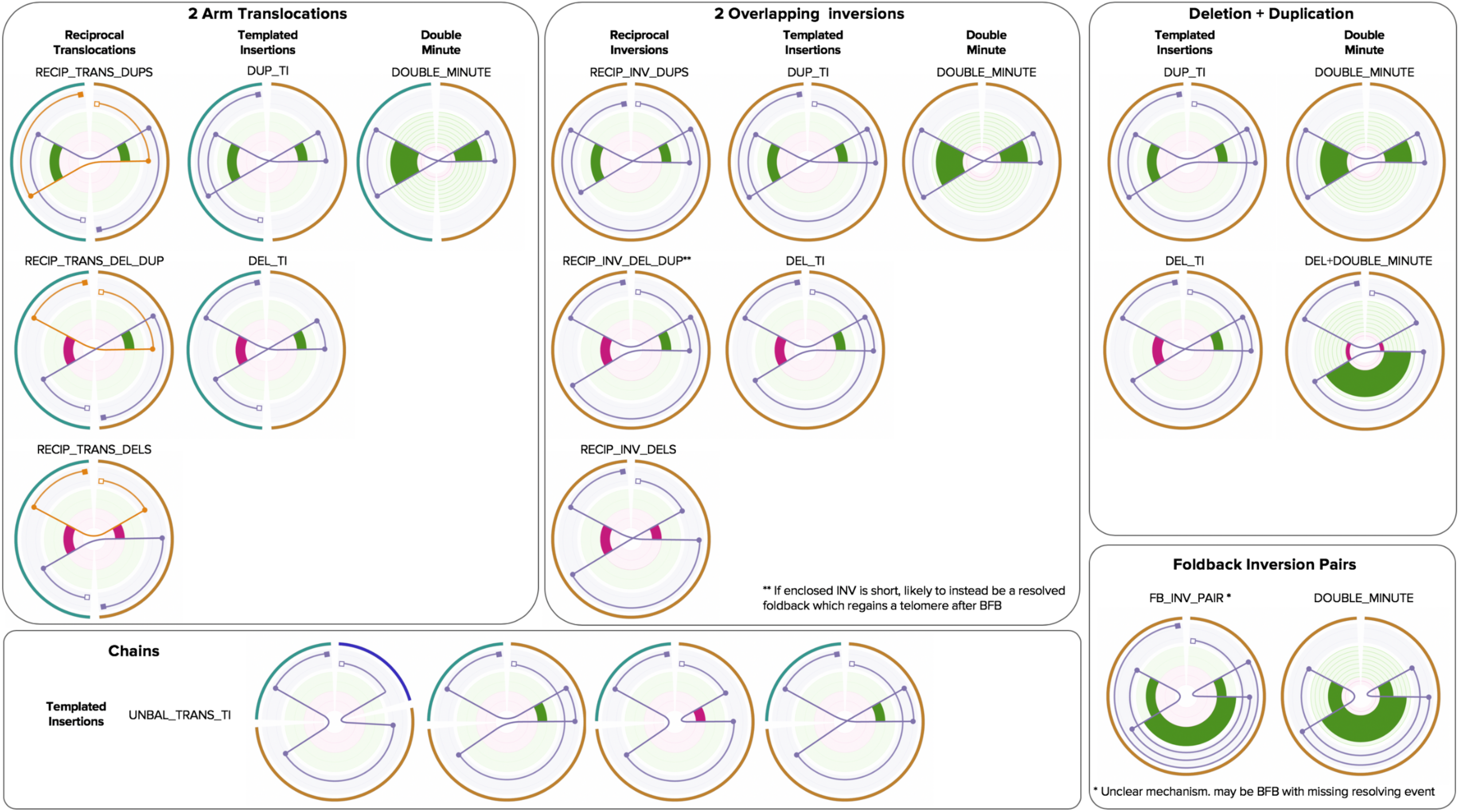
A catalog of consistent 1 break junctions structures in LINX. The lower 3 images show examples of equivalent multiple break junctions events that can be treated as equivalent synthetic versions of the above 1 junction events with a genomic shard (<1000 base segment of templated DNA) inserted.

**EXTENDED FIGURE 5:**
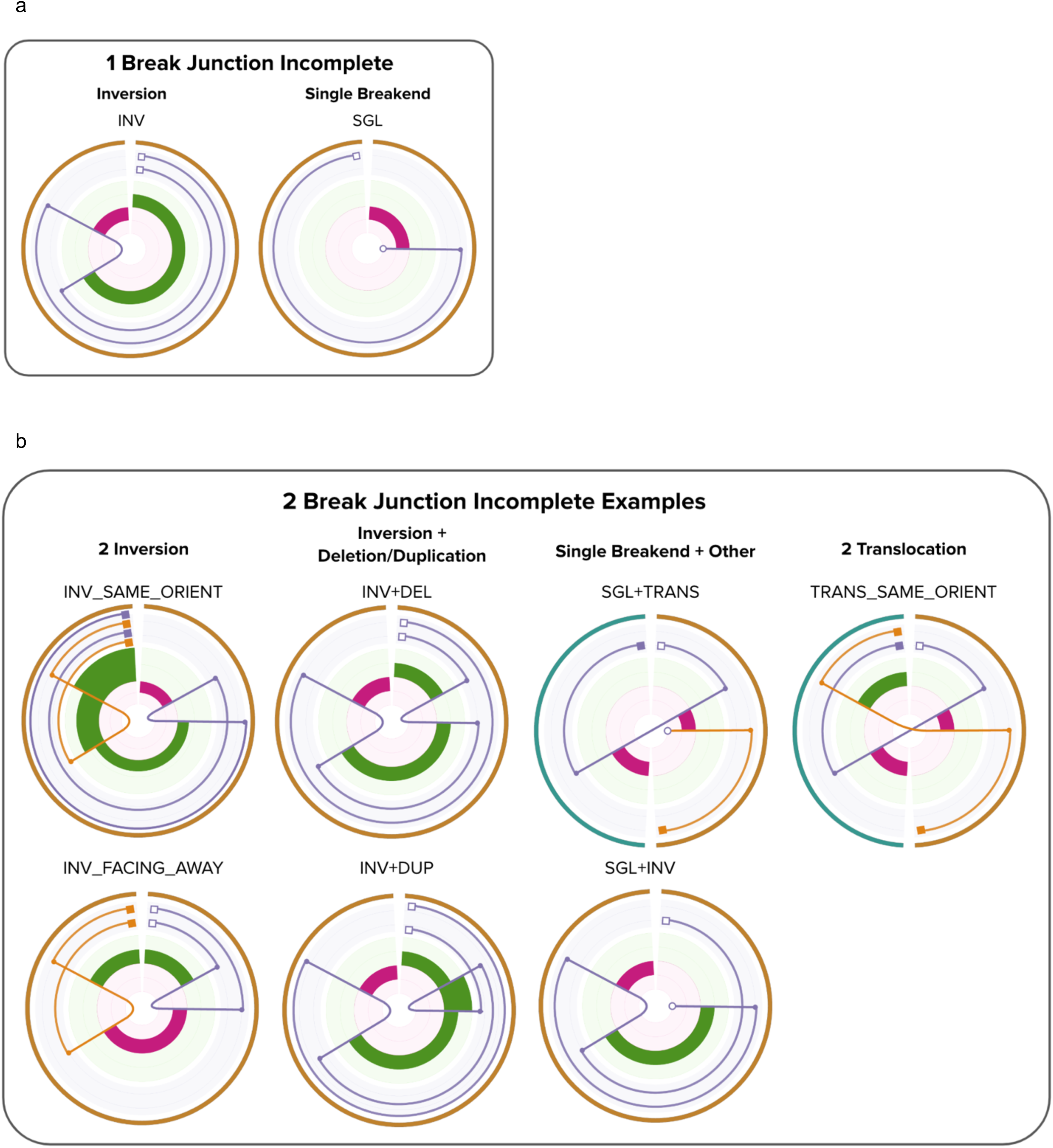
A catalog of consistent 2 break junction structures in LINX. The structures are organised by the geometry of the break junctions.

**EXTENDED FIGURE 6:**
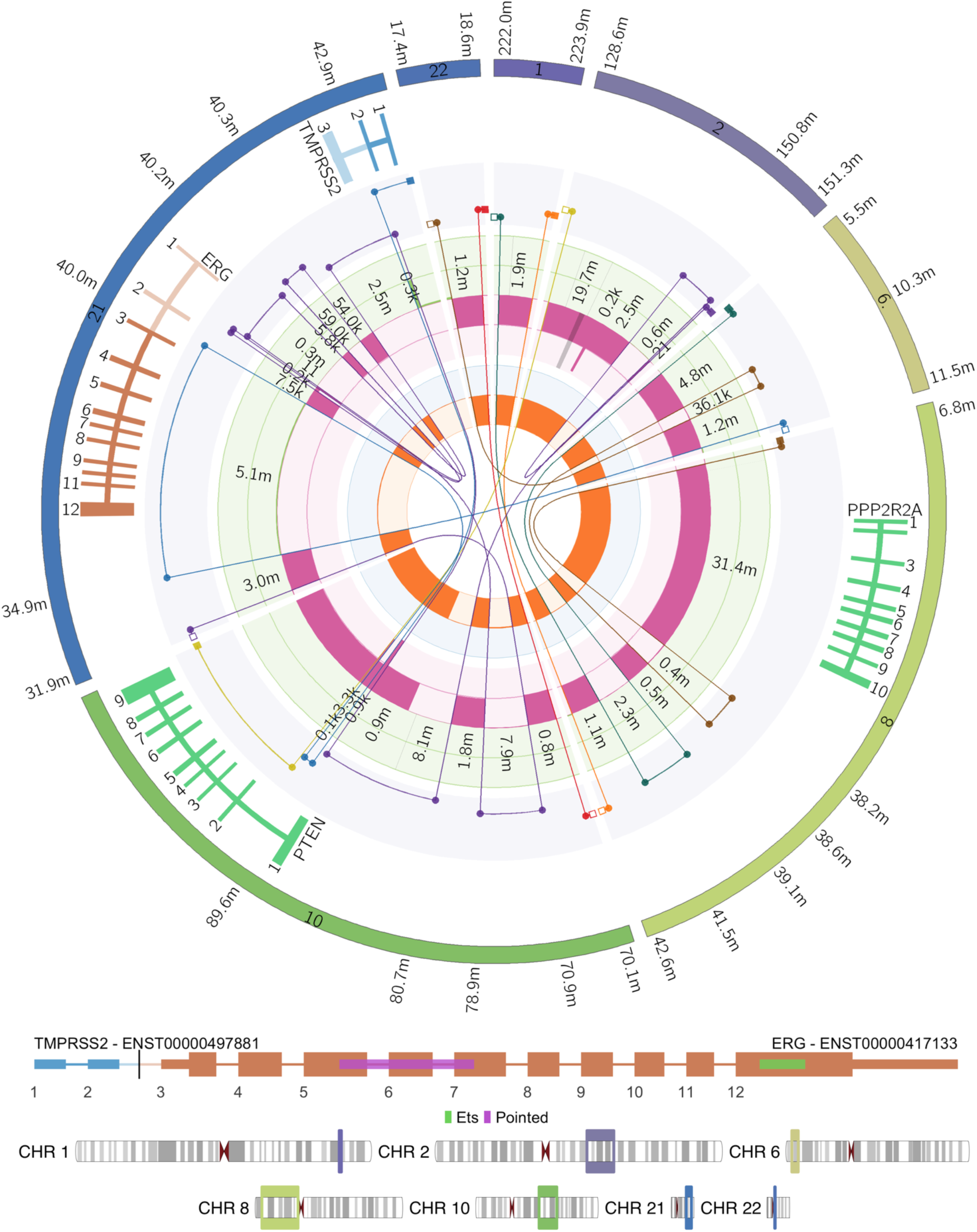
Examples of incomplete 1 and 2 break junction structures in LINX. A cluster is considered incomplete in LINX if it does not create a set of 1 or more consistent derivative chromosomes that each link a telomere and centromere.

## Supplementary Data

**Supplementary table 1**: Cross tool comparison of structural variant calls including validation status for 13 samples

**Supplementary table 2:** Cross tool comparison of somatic copy number and structural variation calling for COLO829

**Supplementary table 3:** Combined STAR-Fusion RNA and LINX DNA fusion predictions for 534 samples

