## Supplementary Methods for "GRIDSS, PURPLE, LINX: Unscrambling the tumor genome via integrated analysis of structural variation and copy number"

### Online Methods

### GRIDSS

GRIDSS version 2 extends the GRIDSS [15] software suite with additional features, tools and capabilities. GRIDSS 2 uses the same genome-wide positional de Bruijn graph break-end assembler used by GRIDSS, as well as the same approach to variant scoring. GRIDSS 2 includes the following additional features, tools and capabilities not present in GRIDSS:

- GRIDSS 2 reports single breakend variants. Single breakends variants are structural variant breakpoints for which only one side can be uniquely mapped. Only single breakend variants supported by an assembly are reported and the assembled sequence is reported as part of the variant call. Soft clipped reads, assembly contigs, and reads with unmapped mates are scored in the same manner as split reads, assembly contigs, and discordant read pairs, but with only a single mapping quality score. Any evidence that supports a specific breakpoint call (split reads, assembly contigs with split read alignments, discordantly mapped read pairs) is considered to support that breakpoint and not either of the corresponding single breakend calls. That is, only assembly contigs with no unambiguous alignment location, soft clipped reads, and read pairs with only one read mapped are considered as providing direct single breakend support. Split read and discordant read pairs can be incorporated into an assembly that provides only single breakend support.
- GRIDSS 2 performs compound realignment of the entire assembly contig. BWA is used to align the entire assembly contig. Assembly contig bases which are soft clipped in the primary alignment reported by BWA are fed back to BWA for realignment. This process is repeated until either either all bases are aligned, or no alignment can be found for the remaining bases. Assembly contigs that do not overlap with the locus of origin of the assembly are filtered out. To ensure that valid assemblies are not unnecessarily filtered, GRIDSS 2 includes both reads of each fragment in the assembly (GRIDSS did not assemble reference-aligning anchoring read when assembling discordant read pairs), and up to 300bp of anchoring reference-supporting sequence is included in the assembled contig. The remaining contigs are treated as split read alignments. To rectify over-alignments in the presence of imperfect breakpoint microhomology, the bounds of each split is adjusted to minimise the edit distance to the reference. Contigs with 3 or more alignment locations are considered to support multiple breakpoints. Crucially, the ability to assemble a contig spanning multiple breakpoints from fragments aligning to a single locus indicates that these breakpoints co-occur on the same chromatid and are thus linked or, in the case of germline variants, phased together.
- Assembly support is tracked per base pair. Fragments are considered to support a breakpoint only if the fragment support spans at least one base pair beyond any breakpoint homology on both sides. This ensures that when a single contig spans both a germline indel and a somatic SV, the fragments originating from the matched normal sample will not be considered as supporting the somatic breakpoint due to their inclusion in the assembly contig as a result of the nearby germline indel, as well as improving variant allele fraction calculations in regions of complex rearrangement.
- Reads alignments containing an insertion or deletion 5bp or larger are included in the analysis. Such reads are treated as split reads with the alignment split into two alignments at the indel location. Indel-containing reads use the same scoring formula as split reads but instead of using the empirical distribution of soft clip lengths for the

library, the distribution of the corresponding insertion or deletion CIGAR element lengths is used.

- Supporting read counts have been replaced by supporting fragment counts. This prevents double-counting of discordantly mapped fragments for which one of the reads contains a split read alignment. A fragment can support a variant either directly through split read, soft clipped read or discordant alignment of a read pair, indirectly through incorporation of one or both of the constituent reads in an assembly supporting the variant, or both directly and indirectly.
- A minimum of two supporting fragments are required to call a variant.
- During *gridss.AllocateEvidence*, reads are preferentially allocated to variants supported by an assembly containing the read.
- GRIDSS 2 supports targeted variant calling on a set of defined regions of the genome. The GRIDSS 2 *gridss.ExtractFullReads* utility is used to extract all reads from fragments overlapping a region of interest. Unlike existing utilities such as samtools [30], *gridss.ExtractFullReads* will extract the mate read and all split read alignments records for all reads overlapping the targeted region regardless of the mate/split alignment location. An example script *example/gridss_targeted.sh* is included in GRIDSS 2 as an example of how to perform targeted calling.
- GRIDSS 2 assembly realignment is performed on the entire assembly contig. Contigs that do not partially align to the originating assembly location are filtered.
- Worse-case assembly performance has been improved by adding an assembly graph path count threshold. Generating 3 assembly contigs with more than 50,000 alternative paths through the assembly graph without advancing the assembly window will flush the assembly window.
- Maximum assembly window size has been reduced by 2.5x and more aggressive assembly read downsampling in high coverage regions is performed.
- The minimum alignment mapping quality score (MAPQ) require for a read to be considered as aligned has been increased to 20, and split reads are only considered if both the primary and supplementary alignments meet the minimum MAPQ.
- The maximum coverage before a region is excluded from analysis has been increased to 25,000x.
- Read alignments containing an insertion of deletion under 5bp are considered consistent with the reference.
- Inversion-like breakpoints are excluded from the minimum variant size. This prevents filtering of foldback inversions.
- When *gridss.SoftClippedToSplitReads* invokes bwa, *-L 0,0* is added to the command line to remove the soft-clipping alignment penalty. This prevents 1bp non-template inserted sequences being over-aligned and reported as clean breakpoints with a flanking SNV.
- Added *gridss.AnnotateUntemplatedSequence* utility. This utility aligns any single breakend sequences or non-template inserted breakpoint sequences to an arbitrary reference genome and add an annotation reporting the alignment location. Integrated viral sequence is identified by aligning to a reference of viral sequences.

GRIDSS 2 provides a full per-sample breakdown of all supporting evidence for each variant through the following VCF INFO and FORMAT fields:

- AS, RAS, CAS: counts of assembly contigs supporting a breakpoint originating locally, from the other side of the breakpoint, and from another location respectively.

CAS assemblies support multiple variants and provide linking information about those variants.

- ASSR, ASRP: total number of split/soft clipped/indel-containing reads, and discordant read pairs/reads with unmapped mate contributing to any breakpoint-supporting assembly contig at the breakpoint location. Note that read/read pairs that are assembled into a contig but whose interval of support does not span the breakpoint are not counted. The interval of support for a read/read pair is defined as the interval between the first and the last contig base for which that read/read pair contributed to the assembly.
- SR, RP, IC: counts of split reads and discordantly mapped read pairs, and indel-containing reads that directly supporting the breakpoint.
- BA: counts of assembly contigs support a single breakend at this position. Such contigs are aligned only to the local breakend with the breakend sequence either aligning ambiguously, or unable to be aligned to the reference genome by bwa.
- BASSR, BASRP: total number of split reads or soft clipped reads, and discordant read pairs or reads contributing to any breakend-supporting assembly contig at the variant location.
- BSC, BUM: counts of soft-clipped reads, and reads with unmapped mates at the variant location
- ASQ, RASQ, CASQ, SRQ, RPQ, IQ, BAQ, BSCQ, BUMQ: corresponding quality score contribution for the supporting evidence.
- QUAL, BQ: total contribution to the breakpoint/breakend quality score.
- BANRP, BANSR, BANRPQ, BANSRQ: counts of read pairs/split reads not supporting this breakpoint but assembled into a contig that does supports this breakpoint and their corresponding assembly quality score contribution.
- REF/REFPAIR: count of reads/read pairs spanning the local variant position that support the reference allele. Only reads/read pairs that span across the breakpoint microhomology interval (if present) are counted.
- VF/BVF: count of unique fragment supporting the breakpoint/breakend.
- RF: count of unique fragments supporting the reference allele.
- CQ: variant quality score prior to evidence prior to evidence reallocation
- BEALN: Potential alignment locations of breakend sequence as determined by *gridss.AnnotateUntemplatedSequence*
- BEID, BEIDL, BEIDH: identifiers of assembly contigs and the corresponding 0-based offset in the assembly contig of the bases corresponding to nominal positions of the local and remote breakends respectively. Single breakend variants do not have a remote breakend, and only breakpoint variants only include breakpoint-supporting assemblies.
- CIPOS: For IMPRECISE variants, CIPOS encodes the interval in which the breakpoint could occur and for precise variants, CIPOS encodes the homology interval.
- CIRPOS: corresponding CIPOS of the remote breakend.
- IHOMPOS: interval of inexact homology. A Smith-Waterman alignment of the breakpoint sequence against the reference sequence is performed at both breakends. The reference and breakpoint sequence are extended 300bp from the on either side with the reference extended an additional 10bp to account for potential indels in the alignment. The homology length is the length that the sequence alignment could be extended from the common sequence into the

breakpoint/reference sequence. Alignments containing a soft clip on the common sequence side are classified as alignment errors and ignored. The SSW library [31] is used for which we implemented a JNI wrapper for. Alignment scored 1, -4, 6, 1 for match, mismatch, gap open, and gap extend respectively which correspond to bwa mem alignment scores.

- SC: CIGAR encoding of the anchoring bases for each at least one read/read pair/assembly is aligned and supports the variant. This is encoded as a CIGAR string with a match for each anchoring base that provides support for the variant call, XNX for the interval over which the breakpoint could occur (due to microhomology or an imprecise call), and a deletion CIGAR element for any intervals over which there is no support (such as a small flanking deletion).
- SB: Strand bias of the reads supporting the variant. 1 indicates that reads would be aligned to the positive strand if the reference was changed to the variant allele. 0 indicates that reads bases would be aligned to the negative strand if the reference was changed to the variant allele. Strand bias is calculated purely from supporting reads and exclude read pair support since these are intrinsically 100% strand bias. Note that reads both directly supporting the variant and supporting via assembly will be double-counted. Both breakpoint and breakend supporting reads are included.
- IMPRECISE, HOMLEN, HOMSEQ, PARID, EVENT, CIEND, END, and SVTYPE fields carry their usual meaning as per the VCF file format specifications.
- BEIDL, BEIDH, BEID encode the position of the all supporting assembly contigs. Assembly contigs supporting multiple variants indicate cis phasing of those variants.

#### Somatic call set generation

By default, GRIDSS is a sensitive caller and reports all putative variants supported by at least two well-mapped reads. To generate a set of high and low confidence somatic call set, a somatic filtering script was created that performs the following steps:

- Adjust the nominal call position of all variants to the centre of any breakpoint microhomology interval.
- “normalSupport”: Filter variants with more than 3% of the supporting reads originating from the normal
- “normalCoverage”: Filter variants with breakend coverage of less than 8 fragments coverage.
- “af”: Filter variants with an allelic fraction of less than 0.5% in the tumor.
- “imprecise”: Filter imprecise variants
- “NO_ASRP”: Filter single breakend variants without an assembly containing at least one discordant read pair
- Filter single breakends with a poly-C or poly-G run at least 16bp in the breakend sequence
- “strand_bias”: Filter deletion or duplication breakpoints under 1000bp with a split read strand bias of 0.95 or greater.
- “homlen”: Filter breakpoints with a microhomology of over 50bp
- “ihomlen”: Filter breakpoint with an inexact microhomology of over 50bp which are not deletion or duplications under 1000bp

- “BPI.Filter.SRSupportZero”: Filter deletion or duplication breakpoints under 1000bp that have no split read support either directly, or through assembly
- “BPI.Filter.PRSupportZero”: Filter breakpoints with no discordant read pair support (either directly, or via assembly) which are not deletion or duplications under 1000bp
- “SRNormalSupport”: Filter deletion or duplication breakpoints under 1000bp that have any direct split read support in the normal
- “small.del.ligation.fp”: Filter 100-800bp deletion breakpoints with an inexact microhomology length of 6bp or greater
- “small.inv.hom.fp”: Filter inversion-like breakpoints 40bp or less that have at least 6bp of microhomology
- “small.replacement.fp”: Filter deletion-like breakpoints under 1000bp whose length of sequence inserted at the breakpoint is within 5bp of the deletion length, except those whose edit distance to the deleted bases is at least 0.5 per base, and less than 0.2 per base to the reverse complement.

Breakpoint variants are filtered if either breakend is filtered.

#### Variant annotation

Variants called were annotated as “PON” based on overlap with a panel of normals (PON). A GRIDSS 2 PON was constructed from the 40x coverage WGS matched normals for 3,972 patients using the *gridss.GeneratePonBedpe* utility. If multiple samples for a patient existed, only the normal for the first sample was included in the PON. Variants were aggregated across samples and the following filters were applied:

- Existing GRIDSS 2 filters were cleared
- All imprecise calls were filtered
- Breakpoint variants with GRIDSS QUAL score less than 75 were filtered
- Single breakend variants with a GRIDSS QUAL score of less least 428 were filtered

A breakpoint BEDPE and single breakend BED file were generated requiring at least two supporting PON samples. A 2bp margin of error was allowed for breakpoint variants. If a breakpoint variant overlapped a PON breakpoint at one breakend and was within 2bp of overlapping on the other side, the bounds of the PON breakpoint was expanded to include the new variant. For breakpoint variants with microhomology, the variant was considered overlapping if any homologous position was overlapping. The generated PON files are available at http://resources.hartwigmedicalfoundation.nl/ as GRIDSS_PON_3792v1.zip

Viral insertions are annotated using *gridss.AnnotateUntemplatedSequence*. Single breakend sequences and non-template inserted sequences that do not have an alignment to the reference genome were aligned to a set of human viral reference sequences. Viral reference sequences were obtained from the virus host database (ftp.genome.jp/pub/db/virushostdb/)[32] and filtered to include only viruses associated with the homo sapiens taxid of 9606. The viral sequences were then masked using RepeatMasker with “-no_is -s -noint -norna -species human” parameters. Generation scripts can be found at https://github.com/hartwigmedical/scripts/tree/master/virus.

- Assembly linking: pairs of breakpoints mutually supported by a common assembly contig were annotated as linked by assembly. For assembly contigs spanning more than 2 breakpoints, each adjacent pair was linked with a unique identifier to enable unambiguous traversal of the breakpoint graph.
- Transitive linking: pairs of precise breakpoint variants for an imprecise transitive breakpoint could be found were annotated as linked by transitive call. Transitive breakpoints were identified by identifying a unique simple path through the breakpoint graph where the start and end positions were within 100bp of the transitive breakpoint position, each segment traversed was at least 20bp in length, at most 4 segment segments traversed, with a total traversed length at most 1,000bp. To prevent exponential runtime in highly rearranged genomes, a breadth first search was performed with at most 100,000 paths and at most 1,000 paths per starting breakpoint were considered.
- Simple inversion annotation: pairs of breakpoint with orientations consistent with a simple inversion were annotated as simple inversions if the matching breakends were within 35bp on both sides, no other simple event annotation could be applied, and fragments supporting the constituent variants differed by at most threefold.
- Templated insertion annotation: breakend/breakpoint and breakend/breakend pairs were annotated as simple templated insertions if the breakends had opposite orientations, were within 35bp, no other simple event annotation could be applied, and fragments supporting the constituent variants differed by at most threefold.
- Reciprocal translocation: breakpoint/breakpoint pairs were annotated as reciprocal translocations if the breakends on both sides had opposite orientations, were within 35bp, no other simple event annotation could be applied, and fragments supporting the constituent variants differed by at most threefold.
- Equivalent: variants were annotated as equivalent if variants had a breakend with 5bp of each other and they shared a common breakend sequence. Breakend sequences were truncated to the length of the shorter sequence and were considered matching when the per-base edit distance between breakend sequences was 0.1 or less. For the purposes of this comparison, the nominal breakend sequence was used for single breakends, and the reference sequence of the partner breakend was used for breakpoint variants. For breakpoint variants, the length of the breakend sequence was the maximum of 20 bases, and the width of the interval over which the fragments supporting the partner breakend had anchoring alignments.

Finally, a quality filter was applied to breakpoint variants with a QUAL score less than 350 and single breakend variants with a QUAL score less than 2000. Variants linked to a variant passing the qual filter other than through equivalence were rescued from the quality filtering and were considered to have passed regardless of the actual variant quality score. For each input file, two output files were generated. A high confidence call set containing calls passing all filters, and a low confidence call set containing all calls except those failing the normal support filter or short events with split read support in the normal.

### PURPLE

#### Input

The PURPLE algorithm relies on the BAF and read depth ratio output from AMBER and COBALT respectively. They should both be run before running PURPLE.

It is also strongly recommended to run PURPLE with a high quality set of somatic SNV and INDEL calls and somatic structural variant calls.

##### COBALT

COBALT determines the read depth ratios of the supplied tumor and reference genomes.

COBALT starts with the raw read counts per 1,000 base window for both normal and tumor samples by counting the number of alignment starts in the respective bam files with a mapping quality score of at least 10 that is neither unmapped, duplicated, secondary, nor supplementary. Windows with a GC content less than 0.2 or greater than 0.6 or with an average mappability below 0.85 are excluded from further analysis.

Next we apply a GC normalization to calculate the read ratios. We divide the read count of each window by the median read count of all windows sharing the same GC content then normalise further to the ratio of the median to mean read count of all windows.

Finally, the reference sample ratios have a further 'diploid' normalization applied to them to remove megabase scale GC biases. This normalization assumes that the median ratio of each 10Mb window (minimum 1Mb readable) should be diploid for autosomes and haploid for sex chromosomes in males in the germline sample.

##### AMBER

AMBER calculates the BAF of the tumor sample by finding heterozygous locations in the reference sample from a panel of 1,344,880 common germline heterozygous SNP loci. The loci were chosen by running the GATK HaplotypeCaller over 1700 germline samples and then selecting all SNP sites which are heterozygous in 800 to 900 of the samples.

To ensure that we only capture heterozygous points, we filter the panel to only loci with allelic frequencies in the reference sample between 40% and 65% and with depth between 50% and 150% of the reference sample genome wide average. Furthermore, we filter any loci with a mapping quality < 1 or base quality < 13. This typically yields 500k-540k heterozygous germline variants per patient.

As part of a contamination check, AMBER also finds sites in the tumor that are homologous in the reference sample using the same panel as above. A sample is considered contaminated if at least 2000 of these sites contain 3 or more reads supporting an alt in the tumor. In this case we model the expected number of non-homologous sites using a poisson distribution and estimate a contamination percent. The result of this is included in the amber QC output file.

For more information on how to run AMBER please refer to the readme.

##### Structural Variant Input VCFs (optional)

Providing a high quality set of structural variant calls to PURPLE allows exact base resolution of copy number changes. An accurate estimation of VAF at each breakend also allows PURPLE to infer copy number changes even across very short segments of the genome where a depth based estimation is inaccurate or impractical. Finally, PURPLE also supports recovery of filtered structural variant calls

For these purposes, PURPLE provides full support and integration with the structural variant caller GRIDSS. GRIDSS can be run directly on tumor and reference BAMs. Alternatively a lightweight version of GRIDSS can be used to re-analyse a set of variant calls and provide additional filtering and accurate VAF estimation.

##### Somatic Variant Input VCF (optional)

A high quality set of somatic SNV and INDEL calls can also improve the accuracy and utility of PURPLE. If provided, the variants are used for enhancing the purity and ploidy fit in 2 ways. Firstly, each solution receives a penalty for the proportion of somatic variants which have implied ploidies that are inconsistent with the minor and major allele ploidy. Secondly, for highly diploid samples, the VAFs of the somatic variants are used directly to calculate a somatic variant implied purity.

For both purposes, accurate VAF estimation is essential thus PURPLE requires the ‘AD’ (Allelic Depth) field in the vcf. High quality filtering of artifacts and false positive calls is also critical to achieving an accurate fit.

#### Algorithm

There are 10 key steps in the PURPLE pipeline described in detail below:

1. Sex determination
2. Segmentation
3. Sample purity and ploidy fitting
4. Copy number smoothing
5. Inferring copy number for regions without read depth
6. Allele specific copy number inferring
7. Recovery of structural variants and filtering of single breakends
8. Identification of germline copy number alterations that are homozygously deleted in the tumor
9. QC Status for the tumor
10. Somatic enrichment

#### 1. Sex determination

We examine both the AMBER and COBALT data to independently determine and validate the sex of the patient. This includes detecting the presence of Klinefelter syndrome: a chromosomal disorder resulting in 2 or more X chromosome in a male and which we have found to affect 0.2% of the male samples in our cohort.

To determine the sex of a sample with AMBER, we examine the number of heterozygous loci in our provided BED file of 1.3 million common germline SNPs on the X chromosome, outside the pseudoautosomal region.   Anything less than 1k heterozygous loci is considered male. A typical female has 12-13k heterozygous loci.

To determine sex using COBALT, we first use the reference ratio to determine the number of copies of the X chromosome. A median X ratio greater than 0.65 is interpreted as 2 copies (note that nearly all female samples are very close to a ratio of 1, but a handful are significantly lower with mosaic X loss). If there is only one copy of the X chromosome the sample is male. Otherwise, we check for the presence of the Y chromosome as determined by at least 1000 data points with a median ratio > 0.05. If the Y chromosome is present (in addition to the 2 copies of the X chromosome), then the sample is male with Klinefelter syndrome. In the absence of the Y chromosome the sample is female.

Finally we compare the AMBER and COBALT sexes. If they are inconsistent we use the COBALT sex and flag the sample has having failed sex validation.

#### 2. Segmentation

We segment the genome into regions of uniform copy number by combining segments generated from the COBALT read ratios for both tumor and reference sample, the BAF points from AMBER, and passing structural variant breakpoints derived from GRIDSS. Read ratios and BAF points are segmented independently using the Bioconductor copynumber package which uses a piecewise constant fit (PCF) algorithm (with custom settings: gamma = 100, k =1). These segment breaks are then combined with the structural variants breaks according to the following rules:

1. Every structural variant break starts a new segment, as does chromosome starts, ends and centromeres.
2. Ratio and BAF segment breaks are only included if they are at least one complete COBALT read depth window away from an existing segment.

If the segments identified by the PCF algorithm are not contiguous, then there remains some uncertainty about the actual start position of the segment. To address this, we use the PCF break as the start position but also include a min and max start position to capture the uncertainty. Segments with SV support are never uncertain.

Once the segments have been established we map our observations to them. In each segment we take the median BAF of the tumor sample and the median read ratio of both the tumor and reference samples. We also record the number of BAF points within the segment as the BAFCount and the number of tumor read depth windows within the segment as the depth window count.

A reference sample copy number status is determined at this stage based on the observed copy number ratio in the reference sample, either ‘DIPLOID’ (0.8<= read depth ratio<=1.2), ‘HETEROZYGOUS_DELETION’ (0.1<=ratio<0.8), ‘HOMOZYGOUS_DELETION’ (ratio<0.1),'AMPLIFICATION'(1.2<ratio<=2.2) or 'NOISE' (ratio>2.2). The purity fitting and smoothing steps below use only the DIPLOID germline segments.

#### 3. Sample purity and ploidy

To estimate purity and sample ploidy, we use a model which considers a matrix of all possible sample purities and ploidies and scores each possible combination on a segment by segment basis, based on a set of principles which aim to choose the most parsimonious solution for the fit.

The specific scoring principles applied are the following:

1. **Penalise sub-clonality**: The major and minor allele of each segment should be close to an integer copy number for clonal solutions. Due to sampling noise, small deviations from integer ploidies will be observed even, but larger deviations require subclonal features and are penalised.
2. **Penalise solutions which deviate from diploid heterozygous copy number**: Loss of heterozygosity or loss of gain of chromosomal segments requires assumptions additional biological complexity, with the complexity increasing for larger deviations from diploid. PURPLE uses an event penalty to explicitly penalise deviations from diploid heterozygous copy number
3. **Penalise solutions with implausible somatic SNV ploidies**: somatic SNVs in principle occur on only one chromatid and should not be found on both alleles. Therefore we penalise solutions where SNV ploidies exceed the major allele copy number.
4. **Weigh segments by count of BAF observations**: Segments are weighted by the count of BAF observations which is treated as a proxy for confidence of BAF and read depth ratio inputs.
5. **Place more weight on segments with higher observed BAF**: segments with lower observed BAFs have more degenerate fits and are weighted less in the fit

For each [sample ploidy, purity] combination we calculate a fit score using the following formula

Fit Score = DeviationPenalty * EventPenaltyMultiplier + SomaticDeviationPenalty

The [sample ploidy, purity] combination with the lowest fit score is selected by PURPLE as the final fit score.

Each of the 3 penalty terms is described in detail in the following sections.

##### Deviation Penalty

The deviation penalty aims to penalise [ploidy|purity] combinations which require extensive sub-clonality to explain the observed copy number pattern.

For each [ploidy|purity] combination tested, an implied major and minor allele copy number is calculated based on the observed BAF and depth ratio. A deviation penalty is then calculated for each segment for both minor and major allele based on the implied ploidies. The function used is designed to explicitly capture a set of intuitive rules relating to biological limitations of cancer genomes, specifically:

- For major allele plody > 1 and minor allele copy number > 0 a deviation penalty applies to penalise solutions which imply subclonality.
    - the penalty depends only on the distance to the nearest integer copy number and varies between a minimum of a small baseline deviation [0.2] and a max of 1.
    - small deviations from an integer don't occur any additional penalty, but once a certain noise level is exceeded the penalty grows rapidly to the maximum penalty reflecting the increasing probability that the observed deviation requires an implied non-integer (subclonal) copy number.
    - The deviation penalty is increased more slowly at lower purities reflecting the increased expected noise. This is implemented by modeling the penalty as a normal distribution with the standard deviation a function of the purity.
    - An additional and increasing penalty multiplier applies for implied major allele copy number < 1. This is intended to capture the relative rarity of large homozygous deletions in cancer genomes, so potential solutions with significant amounts of homozygous deletion are penalised significantly.
- An additional and increasing penalty also applies for implied minor allele copy number < 0 to strongly penalise solutions which imply negative copy number and are biologically implausible.

The following chart illustrates the deviation penalty applied for each of minor and major allele copy number at both 30% and 70% purity.

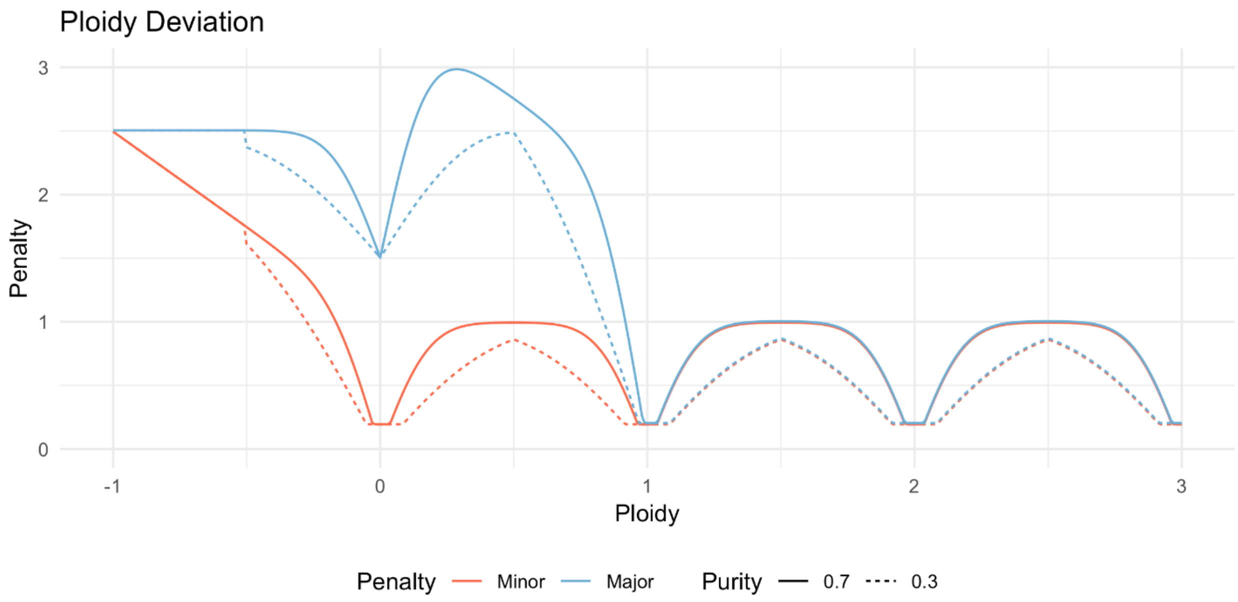

The total deviation penalty is calculated by adding the minor and major allele deviation penalty. It is weighted by the observedBAF (which varies between 0.5 and 1) to reflect that higher BAF regions have fewer possible solutions. The formula used is:

> Deviation Penalty = (MinorAlleleDeviationPenalty + MajorAlleleDeviationPenalty) * ObservedBAF

#### Event Penalty

An event penalty is intended to further penalise [sample ploidy,purity] combinations based on the number of alterations required to get from a normal diploid chromosome to the implied minor and major allele ploidies. In particular, this model penalises higher ploidy solutions that can be highly degenerate and lead to low deviation penalties, but are unlikely to be the most parsimonious or biologically plausible solution.

The event penalty multiplier is given by:

EventPenalty = 1 + 0.3 * min(SingleEventDistance, WholeGenomeDoublingDistance);
WholeGenomeDoublingDistance = 1 + abs(majorAllele - 2) + abs(minorAllele - 2);
SingleEventDistance = abs(majorAllele - 1) + abs(minorAllele - 1);

Note that a diploid segment with implied minor allele copy number = implied major allele copy number = 1 has an event penalty of exactly 1 whilst all other solutions have increasingly higher multipliers as the minor and major allele deviate further from 1. The formula includes an explicit reduced penalty for a doubling of both major and minor allele copy number since there is a known common mechanism of whole genome doubling which can occur in a single event.

The Deviation Penalty and Event Penalty are aggregated independently across all segments that are diploid in the germline and have a tumor depth ratio of <3x the average depth. An average is calculated for each value weighted by the number of BAF observations in each segment. The averaged numbers are multiplied by each other to form an overall ploidy penalty for the sample.

The following chart shows the combined shape of the deviation and event penalty:

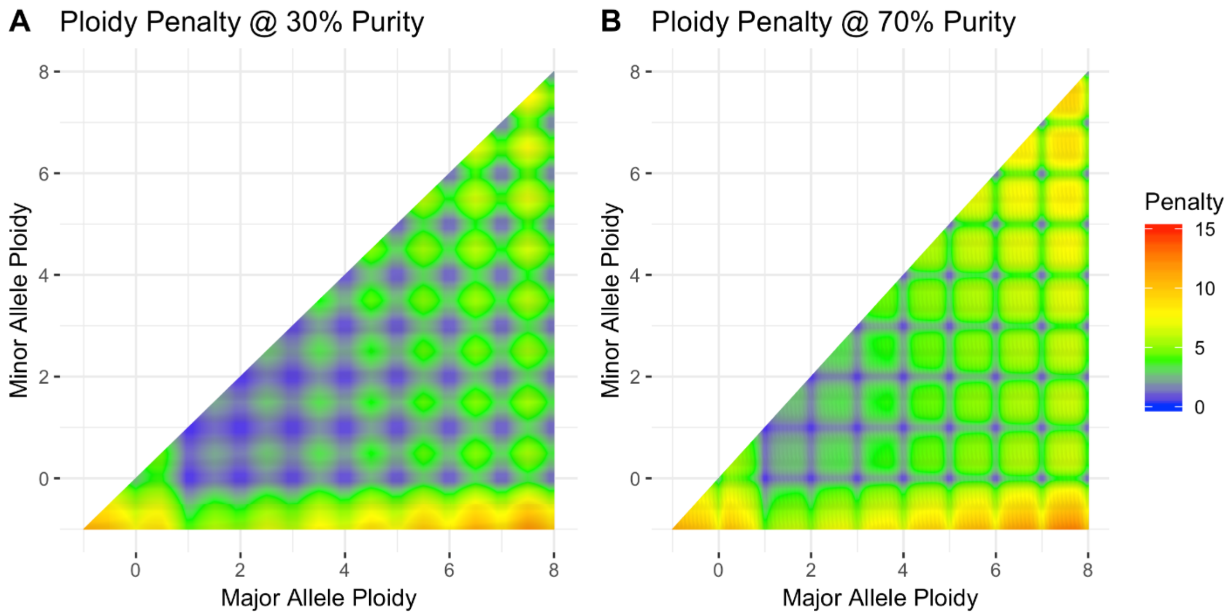

#### Somatic Penalty

If somatic variants are provided, an additional somatic penalty is added to fits which lead to somatic variants with ploidies higher than the major allele copy number, since these are biologically implausible. This feature was introduced primarily to deal with a degeneracy where in certain situations a lower purity, lower sample ploidy solutions may provide a plausible minor and major allele copy number fit to the copy number data, but imply that many SNVs exceed the major allele copy number which is biologically implausible.

The somatic penalty is determined for each [sample plody,purity] combination by sampling 1000 somatic SNV per tumor and comparing the observed ploidy with an upper bound expectation of the variant's ploidy from the 99.9% percentile of a binomial distribution given the major allele at the SNV location. The penalty applied to a single SNV is the max(0,implied SNV ploidy - 99.9% expected bound given the major allele). The somatic penalty is averaged across the 1000 variants and multiplied by a somaticPenaltyWeight [0.3] constant and added to the ploidy penalty.

##### Candidates

While the lowest scoring purity becomes the fitted purity, we also examine other solutions within 10% or 0.0005 of the best solution. These become potential candidates if we need to resort to using the somatic purity described below. We also record the min and max of the purity, ploidy and diploid proportions of the candidates to give some context around the confidence in the solution.

Note that a segment is diploid only if both the major and minor allele are between 0.8 and 1.2 inclusive.

##### Somatic Purity

If any of the candidate solutions are highly diploid (>= 0.97) and there is a wide range (>= 0.15) of valid purities in the candidate solutions we enter somatic mode. Once in this mode, the sample status will be changed from NORMAL to one of HIGHLY_DIPLOID, NO_TUMOR or SOMATIC according to the logic described below.

First we calculate a somatic purity. To do this, we use a kernel density estimator to find significant (n >= 50) somatic variant allele frequency peaks. Each peak implies a tumor purity of twice the frequency. We select the peak that implies the largest purity within the candidate solutions as the somatic purity.

If the somatics are unable to help, either because there are none, or because the somatic purity and the fitted purity are both too low (< 0.17) then we continue to use the fitted purity but flag the solution with a status of HIGHLY_DIPLOID.

If there are only a small number of somatics variants (< 300) with a sufficiently large (>= 0.1) allelic frequency we will flag the solution as NO_TUMOR and use the somatic purity if it exists otherwise fall back on the fitted purity.

If we have not met the criteria for HIGHLY_DIPLOID or NO_TUMOR then we set the status to SOMATIC and use the somatic purity.

#### 4. Copy number smoothing

Since the initial segmentation algorithm is highly sensitive, and there is a significant amount of noise in the read depth in whole genome sequencing, many adjacent segments will have a similar copy number and BAF profile and are unlikely to represent a real somatic copy number change in the tumor. We therefore apply a smoothing algorithm to merge the raw segments into a final set of smoothed copy number regions.

The following criteria apply when deciding to merge segments:

1. Never merge across a segment breakpoint with structural variant support.

2. Do not merge segments if the minimum BAF count in the segments being compared > 0 and the change in observed BAF > 0.03 and the minor allele tolerance is exceeded. The absolute minor allele tolerance is 0.3 + 0.5 * max(copy number of compared segments) / sqrt (min count of BAF points ). For tumors with purity < 20% the absolute tolerances are increased inversely proportional to the purity to allow for greater noise in tumor copy number and BAF measurements.
3. Merge segments where the absolute or relative difference in either the copy number or ref normalised copy number is within tolerances. Absolute copy number tolerance is 0.3 + 2 / sqrt(min depth window count). Relative copy number tolerance is 0.12 + 0.8 / sqrt(min depth window count). Ref normalised copy number uses the actual germline ratios rather than the typical (1 for autosomes, 0.5 for Y etc.). Again, for tumors with purity < 20% the absolute tolerances are increased.
4. Start from most confident germline diploid segment (highest tumor depth window count) and extend outwards in both directions until we reach a segment outside of tolerance. Then move on to the next most confident unsmoothed germline diploid segment.
5. It is possible to merge in (multiple) segments that would otherwise be outside of tolerances if:
    - The total dubious region is sufficiently small (< 30 depth window count or < 50 depth window count if approaching centromere); and
    - The dubious region does not end because of a structural variant; and
    - The dubious region ends at a centromere, telomere or a segment that is within tolerances.

When merging segments, the depth window count (number of COBALT windows) of each segment is used as a proxy for confidence and is used to calculate a weighted average of the copy number. Similarly, the BAF count is used tom calculated the weighted average BAF. The min and max start of the combined region is the minimum from each segment.

Regions that are non-diploid in the germline are typically smoothed over, thus the copy number profile represents the somatic copy number without influence from the germline. However, where smoothing is otherwise not possible PURPLE can sometimes use the reference normalised copy number for this region, ie, the copy number as adjusted for the actual reference read ratio rather than the ideal read ratio. For this to occur, the non-diploid region must:

- Be directly adjacent to a structural variant but not a centromere;
- Not be excessively noisy in the germline (reference read ratio < 2.2x expected);
- Have some depth window support from COBALT;
- be amplified in the tumor relative to the normal, ie, observed tumor ratio > observed normal ratio.

Consecutive non-diploid regions un-interrupted by a structural variant will be merged together.

#### 5. Inferring copy number for regions without read depth information

Where clusters of SVs exist which are closer together than our read depth ratio window resolution of 1,000 bases, the segments in between will not have any copy number information associated with them. We calculate then use ploidies of the structural variants to resolve this.

The outermost segment of any SV cluster will be associated with a structural variant whose ploidy can be determined from the adjacent copy number region and the VAF of the SV. Note that if we are inferring from a lower copy number region into a higher one and the VAF is > 0.75 then we use the read depth rather than the VAF to infer a ploidy. We use the average copy number and read depth of the sample to do this.

Given a SV ploidy, we use orientation of the structural variant to calculate the change in copy number across the SV and hence the copy number of the outermost unknown segment. We repeat this process iteratively and infer the copy number of all regions within a cluster.

When the entire short arm of a chromosome is lacking copy number information (generally on chromosome 13,14,15,21, or 22), the copy number of the long arm is extended to the short arm.

#### 6. Allele specific copy number estimation

Once copy number region smoothing and inference is complete, it is possible there will be regions without BAF points which will result in a unknown allele specific ploidies, since BAF coverage of the genome is limited and many copy number regions can be very small.

For these regions, we infer a BAF and allele specific ploidies of one or more consecutive regions with unknown BAF by first examining neighbouring regions with known allele specific ploidies and inferring based on the observed copy number changes to the unknown regions. Where possible, we assume that only one allele changes copy number in a set of consecutive regions of unknown allele specific copy number. The other allele is held constant.

Which allele copy number to hold remain constant from the neighbouring region depends on a number of rules including on whether the allele specific copy number is known on both sides of the unknown region or only one side.

If only one side of the unknown region is available then we determine which of the major or minor allele of that neighbour to remain constant with the following logic:

- If the unknown region is tiny (<= 30 bases) and is greater in copy number than the neighbour, then hold the minor allele constant.
- Else, find the nearest copy number region where one allele changed by more than 0.5 copy number and hold constant the allele that did not change.
- Failing everything else, keep the minor allele constant.

If there is neighbouring information on both sides of the unknown region, then the following rules apply to determine which allele copy number to hold constant:

- If the minimum copy number of the unknown region is significantly less (> 0.5) than the major allele of both sides, then keep the minor allele constant.
- If both the major and minor allele of the neighbours are significantly different (> 0.5) but the minor allele of one matches the major allele of the other ( < 0.5) then choose the matching allele copy number as the constant copy number.
- Else, if the major allele of the neighbours is significantly different keep the minor allele constant.
- Else, if the minor allele of the neighbours is significantly different keep the major allele constant.

- Else, if the unknown region is tiny (<= 30 bases), and is greater in copy number than the neighbour with the largest number of BAF observations, then hold constant that neighbours minor allele. This rule is intended to deal with the very short overlap which can occur at double strand breaks particularly for LINE insertions.
- Else, if the unknown region is small (<= 1000 bases) and is flanked by large (> 1,000,000 bases) LOH regions, then hold constant the minor allele from the neighbour with the largest number of BAF observations. This rule is intended to ensure that short templated insertions do not break regions of LOH.
- Else, if the unknown region is bounded by a simple DUP with LOH on both sides, keep the minor allele constant.
- Else, find the nearest region on each side (without crossing the centromere) with a change in minor or major allele. If found, re-apply logic from the first three steps.
- Failing everything else, hold constant the minor allele of the neighbour with the largest number of BAF observations.

At this stage we have determined a copy number and minor allele copy number for every base in the genome

#### 7. Structural variant recovery

PURPLE attempts to recover entries from a set of lower confidence structural variants if a recovery vcf is provided. There are two situations where PURPLE will attempt to recover structural variants. The first is when a copy number segment is unsupported by an existing structural variant. The second is to search for a structural variant which could offset the copy number impact of an existing “unbalanced” structural variant break that has a ploidy not supported by the copy number change. A structural variant is considered unbalanced if the unexplained copy number change (ie. the ploidy - copy number change) is greater than 20% of the copy number at the breakpoint and > 0.5. An unbalanced structural variant must also have a min depth window count of 5 in the copy number segments immediately before and after the SV breakpoint. If only one leg of a structural variant is unbalanced but no suitable candidate was found, a single ended breakpoint will be inferred at that position.

Eligible recovery candidates must:

1. Be within 1kb of the min and max range of an unsupported copy number breakpoint or within 1kb of the unbalanced structural variant (if not a single breakend, the other breakpoint must also be within 1 kb of the min-max range of a copy number breakpoint)
2. Not be “AF” filtered in GRIDSS (ie. excluding variants with an allelic fraction of less than 0.5% in the tumor)
3. Have a minimum qual score of 1000 for single breakends and 350 for all others.
4. Have a ploidy of at least 50% of the unexplained copy number change and of at least 0.5.

Following the successful recovery any structural variants we will rerun the segmentation, copy number smoothing and minor allele copy number smoothing with the updated structural variants to produce a final set of copy number segments and breakpoints. Note that the purity estimation does not change.

#### 8. Identify germline copy number alterations that are homozygously deleted in the tumor

During the smoothing process, regions that are homozygously or heterozygously deleted from the germline are smoothed over for the purposes of producing the somatic output. However, as some of these regions are of specific interest, we include them in a separate germline copy number output that contains the homozygous deletes from the germline as well as any deletes that are heterozygous in the germline but homozygous in the tumor.

#### 9. Determine a QC status for the tumor

The status field reflects how we have determined the purity of the sample:

- NORMAL - PURPLE fit the purity using COBALT and AMBER output.
- HIGHLY_DIPLOID - The fitted purity solution is highly diploid (> 95%) with a large range of potential solutions, but somatic variants are unable to help either because they were not supplied or because their implied purity was too low.
- SOMATIC - Somatic variants have improved the otherwise highly diploid solution.
- NO_TUMOR - PURPLE failed to find any aneuploidy and somatic variants were supplied but there were fewer than 300 with observed VAF > 0.1.

PURPLE also provides a qc status that can fail for the following 3 reasons:

- FAIL_SEGMENT - We remove samples with more than 220 copy number segments unsupported at either end by SV breakpoints. This step was added to remove samples with extreme GC bias, with differences in depth of up to or in excess of 10x between high and low GC regions. GC normalisation is unreliable when the corrections are so extreme so we filter.
- FAIL_DELETED_GENES - We fail any sample with more than 280 deleted genes. This QC step was added after observing that in a handful of samples with high MB scale positive GC bias we sometimes systematically underestimate the copy number in high GC regions. This can lead us to incorrectly infer homozygous loss of entire chromosomes, particularly on chromosome 17 and 19.
- FAIL_SEX - If the AMBER and COBALT sex are inconsistent we use the COBALT sex but fail the sample.

#### 10. Somatic enrichment

If a somatic VCF is supplied to PURPLE each variant is enriched with the following fields:

- PURPLE_CN: Purity adjusted copy number surrounding variant location
- PURPLE_MAP: Purity adjusted minor allele copy number surrounding variant location
- PURPLE_AF: Purity adjusted allelic frequency of variant
- PURPLE_PLOIDY: Purity adjusted ploidy of variant
- SUBCL: Subclonal likelihood between 0 and 1
- KT: Forward/reverse kataegis id
- BIALLELIC: Flag to indicate variant is biallelic

More detailed descriptions of the kataegis, clonality and biallelic status are found below.

##### Kataegis

Somatic variants of type C>T and C>G in a TpCpN context are annotated as showing Kataegis [33] if there are three or more mutations of that type, strand and context localised within a region with an average inter-mutation distance of <= 1kb. The annotation describes the strand it was found on along with an id that is shared by grouped variants.

##### Clonality and Biallelic status

For each point mutation we determined the clonality and biallelic status by comparing the estimated ploidy of the variant to the local copy number at the exact base of the variant. The ploidy of each variant is calculated by adjusting the observed VAF by the purity and then multiplying by the local copy number to work out the absolute number of chromatids that contain the variant.

We mark a mutation as biallelic (i.e. no wild type remaining) if Variant Ploidy > Local Copy Number - 0.5. The 0.5 tolerance is used to allow for the binomial distribution of VAF measurements for each variant. For example, if the local copy number is 2 than any somatic variant with measured ploidy > 1.5 is marked as biallelic.

For each variant we also determine a probability that it is subclonal. This is achieved via a two-step process.

First, we fit the somatic ploidies for each sample into a set of clonal and subclonal peaks.

We apply an iterative algorithm to find peaks in the ploidy distribution:

- Determine the peak by finding the highest density of variants within +/- 0.1 of every 0.01 ploidy bucket.
- Sample the variants within a 0.05 ploidy range around the peak.
- For each sampled variant, use a binomial distribution to estimate the likelihood that the variant would appear in all other 0.05 ploidy buckets.
- Sum the expected variants from the peak across all ploidy buckets and subtract from the distribution.
- Repeat the process with the next peak

This process yields a set of ploidy peaks, each with a ploidy and a total density (i.e. count of variants). To avoid overfitting small amounts of noise in the distribution, we filter out any peaks that account for less than 40% of the variants in the ploidy bucket at the peak itself. After this filtering we scale the fitted peaks by a constant so that the sum of fitted peaks = the total variant count of the sample. We mark a peak as subclonal if the peak ploidy < 0.85.

Secondly, we can calculate the subclonal likelihood for any individual variant as the proportion of subclonal variants at that same ploidy.

### LINX

LINX is an annotation, interpretation and visualisation tool for somatic structural variants using genome-wide GRIDSS breakpoint and PURPLE copy number data as input. The primary function of LINX is grouping together individual SV calls into distinct events and properly classify and annotating the event to understand both its mechanism and genomic impact.

#### Key concepts in LINX

##### LINX terminology and conventions for linking proximate breakends

###### Assembled vs inferred links

In LINX, links are chromosomal segments connected by break junctions which are predicted to form part of a derivative chromosome. Assembled links are those that were linked by in a single assembly by GRIDSS as being and are very high confidence somatically phased. All other links are inferred, based on proximity, topology and copy number characteristics using the chaining logic described below.

###### Templated insertions

We have adopted the term ‘templated insertion’ as has been used previously [34] to describe any piece of DNA which is a templated sequence from a section of the ref genome flanked by breakends on either side inserted elsewhere (either locally or on a remote chromosome) into a chain to form part of a derivative chromosome.

##### ‘Shards’ and ‘synthetic’ events

A special and very common case of templated insertions we observe are very small templated genomic fragments of up to several hundred bases in length, which are frequently inserted into breakpoints without disruption at the source site for the inserted sequence. These have been observed previously [35] and termed as genomic ‘shards’. In LINX we model shards explicitly as short templated insertion lengths of less than 1k bases. These inserted sequences can make simple events such as deletions and tandem deletions appear to have complex geometries. For example, if we have a simple short deletion with a shard inserted, and the templated sequence of the shard is from another chromosome the deletion now presents notionally as a chained pair of translocations. Where more than 1 shard is inserted, the complexity can grow even further. LINX simplifies events that could be explained as a 1 or 2 break cluster and where shards which do not overlap the region where they are inserted and marks them as the simple type and marks the cluster as ‘synthetic’.

Extended Figure 3 shows a number of examples of synthetic events with the shards marked.

###### Deletion bridges, anchor distance & overlapping deletion bridges

We use the term ‘deletion bridge’ as defined previously [12] to refer to sections of DNA loss between 2 breakpoints on the same chromatid that are fused to other segments of the genome.
GRIDSS provides an anchor support distance for each structural variant breakend which is the number of bases mapped to the reference genome at that breakend as part of the assembly contig, which is typically in a range from 29 bases (the minimum anchor distance for GRIDSS to be able to call) up to approximately 800 bases for short read sequencing. Any other breakend that falls within this anchor distance cannot be ‘cis’ phased with the variant as the contig was able to be mapped past the breakend and the 2 breakends are ‘trans’. Trans breakends within this distance are common (see figure 2f). One possibility is that the

breakends could occur on the other chromatid, but this highly unlikely as there is no reason to expect 2 different chromatids to both be damaged within a few hundred base region. Much more likely is that when the double stranded break occurred, that there was significant overlap between the break locations on the 2 strands and the overlapping break ends have been replicated prior to fusing with other genome. This is highly analogous to a deletion bridge except with small sections of replication of DNA instead of loss. LINX uses the term ‘overlapping deletion bridge’ to describe this break topology.

#### Overview of event classification system in LINX

LINX attempts to classify all variants into a set of consistent events, i.e. events that transform the genome from one stable configuration into another. The 4 basic categories of events that we have are simple deletions, tandem duplications, insertions and two break events (including both reciprocal events and unbalanced translocations). For events with 3 or more variants that cannot be simplified into the above types, the number of possible configurations quickly explodes so we group these together in COMPLEX and analyse them via topological features rather than as distinct classes.

A key assumption in LINX is that each derivative chromosome arm in a stable configuration must connect a telomere to a centromere (since centromere to centromere joins will cause unstable breakage fusion bridge and telomere to telomere joins will have no centromere and will be ultimately lost during stochastic mitosis processes). A special case is allowed in highly restricted circumstances for double minute chromosomes which are circular and have no telomere or centromere but are highly positively selected for. This assumption means that variants such as a lone head to head or tail to tail inversion are considered incomplete, and in these cases we intensively search for other variants which may have occurred concurrently and could restore a stable configuration. Because of limitations of both input data accuracy and completeness and our clustering and chaining algorithm, many COMPLEX clusters will not be fully resolved to a stable configuration although it is assumed that such a resolution exists. Furthermore, we have a number of residual 1 and 2 clusters (eg. a lone inversion) which are inconsistent cannot be accurately clustered and hence we classify them as INCOMPLETE.

Ultimately we classify each cluster into 1 of 7 major event super types:

| Event Super Type | Description |
|---|---|
| SIMPLE | Single break junction cluster which forms a local deletion, tandem duplication or unbalanced translocation |
| RECIPROCAL | Reciprocal inversion or translocation events forming from 2 concurrent breaks interacting with each other |
| TEMPLATED INSERTION | DEL or DUP or unbalanced translocation (‘chain’) with templated insertion |
| INSERTION | SV that are formed by the insertion of a templated piece of DNA normally via either a mobile line element or virus. |
| DOUBLE_MIUNTE | Any cluster where all variants form part of a double minute |
| COMPLEX | Clusters with 3 or more variants that cannot be resolved into one of the above categories |

| | |
|---|---|
| INCOMPLETE | 1 or 2 breakpoint clusters which are inconsistent, but cannot be clustered further OR clusters which are inferred from copy number changes only |

A brief overview of the non single break junction events types is given below:

Reciprocal events

Linx models reciprocal 2-break junction events as events that could be caused by the interaction of 2 simple local concurrent breaks which would normally form deletes and tandem duplications. Depending on whether the breaks are on the same or opposite chromosomes this forms reciprocal translocations or reciprocal inversions. Note that in the translocation case, if one side of the reciprocal event is subsequently lost either before or after repair, then we observe an unbalanced translocation.

The possible geometries for reciprocal events supported by LINX are explained in the table below and drawn in extended figure 4:

| Interacting Break Types | Same chromosome (Inversion) | Translocation |
|---|---|---|
| Concurrent double stranded breaks | **RECIP_INV** - 2 facing inversions with outer breakends overlapping | **RECIP_TRANS** - 2 translocations forming deletion bridges on both arms. |
| Concurrent tandem duplications | **RECIP_INV_DUPS** - 2 facing inversion with inner breakends overlapping | **RECIP_TRANS_DUPS** - 2 translocations with facing breakends on both arms |
| Tandem Duplication + Double Stranded Break | **RECIP_INV_DEL_DUP -** inversion enclosing inversion with opposite orientation | **RECIP_TRANS_DEL_DUP -** 2 translocations forming a deletion bridge on one arm and and facing breakends on other arm |

A facing pair of foldback inversions (FB_INV_PAIR) is also classified as a reciprocal, although the mechanism for forming this structure is unclear. It is possible that many of these events are formed from a breakage fusion bridge event but have not been properly clustered with a resolving break junction which may be distant in a breakage fusion bridge scenario.

Templated insertions

For the 4 reciprocal event cases above involving duplication (ie. RECIP_INV_DUPS, RECIP_INV_DEL_DUP, RECIP_TRANS_DUPS & RECIP_TRANS_DEL_DUP), the same break junctions can be alternately chained to form a single derivative chromosome with a templated insertion (see extended figure 4). LINX gives precedence to the reciprocal interpretation, but if any of the duplicated segments bound a telomeric or centromeric loss of heterozygosity, the reciprocal interpretation is implausible

A deletion and duplication can together also form either a duplication or deletion with templated insertion structure (extended figure 4) identical to the 2 inversion case but with the inserted segment in the opposite orientation. Unlike inversions, simple deletions and tandem duplications are consistent standalone events and are common genomic events so some of these structures may be clustered incorrectly where separate DEL and DUP events are highly proximate or overlapping by chance

##### Insertions

An insertion event is modelled by LINX as a pair of structural variants which inserts a section of templated sequence from either another part of the genome WITHOUT disruption to the chromatid at the source location OR from an external sequence such as an insertion from a viral genome.

The most common class of insertion in tumor genomes by far are mobile line element insertions, which are not typically active in the germline, but can be highly deregulated in many different types of cancer. Mobile line elements frequently insert short sequences of their own DNA sequence and templated segments from adjacent to the source line element, with sometimes many segments from the same source location being inserted at multiple locations around the genome [36]. Mobile LINE elements can also cause SINE and pseudogene insertions. LINE insertions breakpoints can be often difficult to map correctly on both ends, since they typically involve a repetitive LINE element at the start of the insertion element and a poly-A section at the end of the inserted section. LINX uses a combination of previously known line element source information and identification of both the local breakpoint structure and POLY-A sequences to classify both fully and partially mapped breakpoints as LINE insertions.

##### Double minute

Any cluster which is predicted to form a closed loop by LINX without a centromere is resolved as a double minute. All variants must form part of the double minute to be classified as event type double minute, although double minutes may also occur as part of a complex cluster. An exception is made for a simple DUP double minute clustered with an enclosing DEL, which is classified as double minute despite the DEL not being a part of the double minute structure.

##### Complex events

'Complex' events are simply clusters with 3 or more variants or 2 clusters that can be fully chained, that cannot be resolved into either a simple or synthetic type of insertion, del, dup or 2-break event.

Complex events may be formed by any combination of non-mutually exclusive processes including multiple concurrent breaks, replication prior to repair, breakage fusion bridge processes. Local topology annotations in LINX are intended to shed light on these complex processes.

##### Incomplete & inferred events

There are a number of possible configurations which are not 'COMPLEX' by the above definition since they are formed from 1 or 2 SVs, but lead to inconsistent genomes or involve single breakends. For these clusters there is assumed to be missing SVs, potential false positive artefacts or under clustering and they are marked as INCOMPLETE.

INCOMPLETE includes but is not limited to the following configurations:

- Lone inversion
- Lone single breakend
- Lone inferred breakend
- Any 2 cluster with a single or inferred breakend that cannot be resolved as LINE or inferred as a synthetic.
- Any 2 cluster which cannot be chained OR resolved as either a LINE, synthetic or reciprocal event

Clusters of 2 inferred breakends are also classified in this category. Many of these are likely artefacts due to residual large scale GC biases affecting coverage in our sequencing.

### LINX Algorithm

There are 4 key steps in the LINX algorithm:

- Annotation of genomic properties and features
- Clustering of SVs into events
- Chaining of derivative chromosomes
- Gene impact and fusion prediction

The following schematic outlines the overall workflow in the LINX algorithm. Each step is described in detail below.

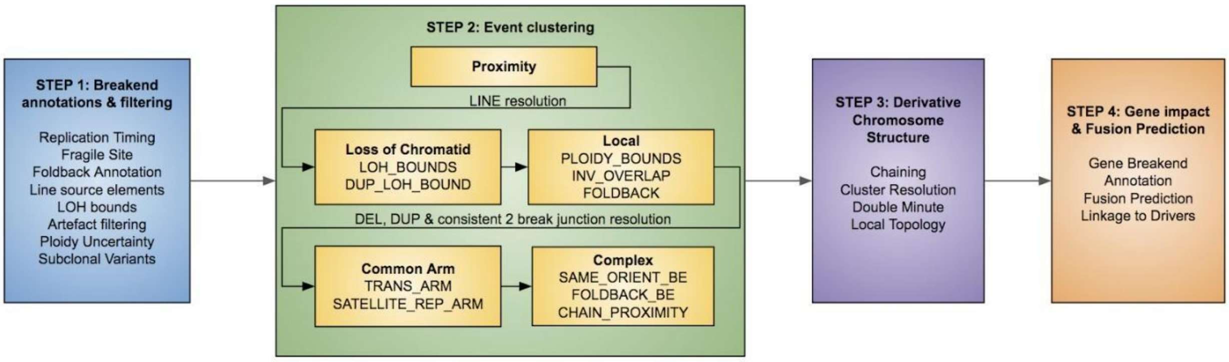

#### 1. Annotation of genomic properties and features

To help resolve and characterise events, LINX first annotates a number of genomic properties.

##### Externally sourced genomic annotations

Each breakend is first annotated with the following information from external sources

- Whether it is in a known fragile site [19]
- Whether it is in a known LINE source element [21]
- The replication timing of the breakend [36]

##### Identification of foldback inversions

Foldback inversions are important structural features in the genome since they are a hallmark of the breakage fusion bridge process and are used by LINX in a number of ways in both the clustering and chaining algorithms. They can also be objectively identified independently of the clustering and chaining so it is useful to identify them upfront. We perform a genome wide search for foldback inversions. A pair of breakends are marked as forming a foldback if they meet the following criteria:

- the breakend orientations are the same and are consecutive (ignoring any fully assembled interim breakends) and after allowing for overlapping deletion bridges on both ends, specifically both:
    - The outer breakend may be overlapped by a variant within it's anchor distance
    - The inner breakend may not have a facing breakend within it's anchor distance

- the breakends belong to a single inversion or are linked by an assembled or short chain (<= 5K bases)
- A single breakend where the other end of the structural variant is assembled to itself via a chain
- Neither breakend forming the foldback is linked via assembly to another breakend.

Identification of suspected LINE source elements

LINE source elements are also important genomic features and are modelled in LINX as regions of ~5000 bases which we suspected are the source for templated LINE insertions. LINE insertion type SVs are common in cancer genomes and typically present as a pair of balanced SVs representing a templated sequence from around the source element with a poly-A tail inserted into random locations in the genome although favoring a A|TTTTT motif [21] for the insertion site with no net copy number change at either source or insertion site. However, due to both the repetitive nature of the LINE source elements and the difficulty of accurately sequencing across the poly-A tail, one or both of the SVs that make up the insertion may be mapped as a single breakend (failing to uniquely map on the other side) OR be missed altogether. The lone remaining breakend can easily be mistaken as an unbalanced translocation so it is important to correctly identify as a LINE element. This picture can be complicated even further by the fact that many overlapping fragments from single source location may be copied to many different locations in the same genome, each potentially with one or both sides incompletely mapped. This can appear as a very complex feature if not broken into the constituent insertions that it represents.

Although we already annotate 123 well ‘known’ mobile line source elements which have been previously discovered, there are many more potential mobile line source elements and many are highly sample specific. We look exhaustively for likely LINE source elements in each individual tumor genome. Specifically each breakend may be classified as being a ‘Suspected’ line source element if:

- there are 2+ breakends within 5kb which both have a poly-A insertion sequence
- OR there are 2+ translocations within 5kb with at least one not forming a deletion bridge of < 30 bases AND at least one SV within 5kb having an INS sequence containing at least 11 repeated As or Ts.
- OR we find at least 1 translocation with a remote single breakend forming a deletion bridge of < 30 bases AND EITHER at least one SV also within 5kb OR the remote single breakend HAVING an INS sequence containing at least 11 repeated As or Ts.

The suspected LINE source element is also checked that it is not a potential pseudogene insertion by checking that there is no deletion within 5kb of the suspected source element that matches an exon boundary at both ends.

Identification of LOH boundaries

We also identify each pair of breakends flanking regions of Loss of Heterozygosity (LOH), restricted to cases where there is no subset of the region with homozygous loss that is not caused by anything other than a simple deletion. This is a useful annotation as since an entire chromatid is lost for the whole distance between these 2 breakends (and since there is no homozygous loss we know it is the same chromatid lost at both ends, not two overlapping losses) then the structural variants are very likely to have occurred at the same time.

Note that an uninterrupted deletion or tandem duplication cannot theoretically form an LOH boundary with another variant and hence these are excluded from LOH boundaries.

Long DEL and DUP length calculation

Shorter deletions and tandem duplications are found frequently as standalone events in the tumor genome, but longer standalone deletions and duplications are relatively rare and when they do occur are often associated with more complex events. The following method is used to determine a characteristic length threshold for each sample for what is considered a 'long' DEL or DUP.

- find all DUPs and DELs on arms with no inversions (inversions are used as a proxy for the presence of complex events)
- Set LONG_DUP_LENGTH to the length of the longest DUP excluding the 5 longest DUP lengths (normalised for the proportion of arms without inversions). Min =100k, Max = 5M
- Set LONG_DEL_LENGTH to the length of the longest DEL excluding the 5 longest DEL lengths (normalised for the proportion of arms without inversions). Min =100k, Max = 5M

The threshold is subsequently used in clustering rules by LINX.

Artefact filtering

Prior to clustering and chaining, LINX applies additional artefact filtering. This is due to the fact that depending on sequencing quality, we may still find a number of false positive artefacts in our data. False positive artefacts are typically either SVs with little or no copy number support (ie copy number change at both breakends < 0.5) or inferred SV breakends from the copy number analysis. Unfortunately these can be difficult to distinguish from bonafide subclonal variants with ploidies of 0.5, and from genuine clonal variants where we have missed an offsetting SV call which netted out the copy number change.

To remove residual artefacts, but preserve genuine subclonal variants, we limit filtering to 5 very specific situations which strongly appear to be artifactual in our data:

- **Equivalent breakend calls -** Any single breakend marked as 'eqv' in GRIDSS assembly data or which matches the orientation and the breakend of another SV within 1 base is filtered prior to clustering
- **Spanning SV calls** - if an SV's breakends match orientation and position (to within 1 base) of the breakends of 2 other SVs, and these other 2 SVs are assembled into a templated insertion on their other breakends, then the spanning SV is considered an artefact.
- **Short foldbacks Inversions (<100 bases) unsupported by copy number change at both breakends**- Typically foldback inversions range from several hundred to several thousand bases. However, we also frequently find many very short foldbacks inversions in highly damaged samples. Across the cohort as a whole we find these to be overwhelmingly low ploidy and with little copy number support. Hence we mark foldback inversions <100 bases in length as artefacts if both start and end copy number change < 0.5 or VAF < 0.05 at both ends.
- **Isolated translocations and single breakends unsupported by copy number change at both breakends -** These also common artefacts and similar to foldback inversions, we find a much higher rate of low ploidy variant calls unsupported by copy number. We filter if both breakends of a translocation are > 5000 bases from another variant AND both breakends have copy number change <0.5 AND the insert sequence is not polyA (which may indicate a Line insertion - see below).
- **Neighbouring inferred breakends with opposite orientation and matching or overlapping ploidy** - Residual GC bias and other forms of copy number noise can cause many inferred segments to be called. These are unhelpful in chaining and are resolved in pairs of offsetting variants as artefacts

All of the filtered variants will be marked as resolved type = ARTEFACT and will be restricted from any subsequent clustering.

Estimation of ploidy and ploidy uncertainty per SV

We know that by definition ploidy must equal the copy number change at start breakend and the copy number change at end breakend for each and every structural variant. However, often times the ploidy and copy number start and end may not match up for one of several reasons including measurement error, false positive SV artefacts and false negative SV calls adjacent to one or both of the breakends for the structural variant.

To allow for accurate chaining we would like to have a single consolidated ploidy estimation and an idea about the uncertainty in the ploidy for each SV. Since the distribution of the errors in our measurements are unknown and fat tailed due to potentially missing and false positive data, we create a simple model for a reasonable estimation of the likely ploidy range.

We use 3 steps in the process:

*1. Estimate an uncertainty for copy number change at each breakend*

For this use the principle that the uncertainty in copy number change is driver primarily by the uncertainty in the copy number of the least confident adjacent copy number region which in turn is driven primarily by the number or read depth windows used to estimate the length of the adjacent regions.

Hence we use the following formula to calculate a copy number uncertainty

> CNChangeUncertainty = MAX(maxAdjacentCopyNumber * BaseRelativeUncertainy [0.1],BaseAbsoluteUncertainty [0.15])+ MAX(AdditionalAbsoluteUncertainty [0.4],AdditionalRelativeUncertainty [0.15]*maxAdjacentCopyNumber)/SQRT(max(minAdjacentDepthWindowCount,0.1))

If the mindAdjacentDepthWindowCount = 0, then this means the segment is inferred by the SV ploidy in PURPLE already and no copy number estimate is calculated.

For the special case of foldback inversions, if the flanking depth window counts are both higher than the internal depth window count, a single copy number change observation of half the combined copy number change is made with the confidences determined from the flanking windows

*2. Estimate an uncertainty for raw ploidy*

The raw ploidy of the SV is already estimated in PURPLE by multiplying the purity adjusted VAF of the SV by the copyNumber at each breakend. The VAF estimate depends ultimately on the measured readcount of supporting tumor fragments which is a binomial distribution.

To estimate the uncertainty in the VAF, we estimate the 0.5% and 99.5% confidence intervals of the true read count from the observed read count and then calculate the ploidy uncertainty as half the relative range of the confidence interval. We also add half the minimum of the 2 breakend copy number uncertainties to reflect the copyNumber impact on uncertainty. This gives the formula:

Ploidy Uncertainty = Ploidy * (ReadCountUpperCI-ReadCountLowerCI) / 2 / ObseverdReadCount + 0.5 * min(CNChangeUncertaintyStart,CNChangeUncertaintyEnd)

*3. Average the 3 ploidy predictions and estimate a consolidated uncertainty*

Weight the observations by the inverse square of their estimated uncertainties:

consolidatedPloidy = SUM[Observation(i)*(1/Uncertainty(i)^2)] / Sum[1/Uncertainty(i)^2]

The combined uncertainty is estimated as the square root of the weighted sum of squares of the difference between the final ploidy estimate and each individual estimate, but capped at a minimum of half the input uncertainty. Ie.

consolidatedUncertainty = SQRT( countObservations / (countObervations-1) * SUM[1/Uncertainty(i)^2*(MAX(Observation(i)-consolidatedPloidy,Uncertainty(i)/2))^2] / Sum[1/Uncertainty(i)^2] )

##### Identification of subclonal variants

We identify high confidence subclonal variants using the uncertainty bounds, using the threshold of maximum ploidy < 0.75. Clonal and subclonal variants are unlikely to have occured at the same time and hence all subclonal variants are excluded from clustering with any variant that does not overlap in ploidy uncertainty and does not have a ploidy within 0.5 of the subclonal variants. Proximity clustering is still allowed, since the ploidy estimates for proximate variants are more uncertain.

#### 2. Clustering of structural variants into events

LINX uses a clustering routine to classify events. All SVs within a sample are grouped into clusters in 5 steps

- Proximity clustering
- Resolution of LINE clusters
- LOH & local overlap clustering
- Resolution of simple events
- Advanced clustering rules

##### Proximity clustering

Any SV with a breakend within the specified proximity_distance (defaults to 5K bases) of another SV's breakend causes the SVs to be clustered. An exception is made for overlapping DELs, which are split out into separate clusters, on the assumption that overlapping DELS must be on different chromatids.

##### Resolution of LINE clusters

Line insertions frequently involve repetitive sequences, and are often only partially resolved. We resolve a cluster as type LINE if:

- It contains a suspected LINE element AND (the cluster has <=10 variants OR at least 50% of the SVs in the cluster have a known or suspected breakend)

- OR if every variant in the cluster touches a KNOWN line element AND at LEAST one of the variants is a translocation
- If the cluster has 2 or less variants both of which are single or inferred breakends and at least one PolyA insert sequence

LINE clusters are excluded from all subsequent clustering rules.

Loss of chromatid bounds

*Loss of chromatid bounds*

The 2 breakends forming the bounds of any LOH region not disrupted by a homozygous deletion are clustered together, reflecting the fact that both ends of the lost chromatid must have been lost at the same time.

For LOH regions which are disrupted by 1 or more homozygous deletions, both chromatids are presumed to have been deleted in separate events and one deletion may enclose the other deletion OR they may overlap each other. In the enclosing case, for homozygous deletions which are in a region where the surrounding LOH bounds wither form a simple DEL or are already linked or extend to at least the full arm, then we cluster the 2 homozygous deletion bounds. Conversely if all the homozygous deletions inside a LOH region are simple DEL or are already linked then we can cluster the 2 LOH region bounds. Finally, in the overlapping case, if all the overlapping deletion bounds are from simple DELs or are already linked except for 2 breakends, then link the remaining 2 breakends.

Note that variants with breakends that bound an LOH are not permitted to link with breakends in the bounded LOH region via any subsequent rule, since by definition they are expected to be on the other chromatid.

*Chaining bounds for DUP variants causing LOH*

No breakend in a cluster can chain across an LOH which has been caused by a breakend in the same cluster. Hence if the other breakend of a DUP type variant bounding an LOH can only chain to only one available (not assembled, not LINE) breakend prior to the LOH, then we cluster the DUP and the other breakend.

Local Clustering

*Major allele copy number bounds*

The major allele copy number of a segment is the maximum copy number any derivative chromosome which includes that segment can have. Hence a breakend cannot chain completely across a region with major allele copy number < ploidy of the breakend, or partially across the region with a chain of ploidy more than the major allele.

Therefore any breakend is clustered with the next 1 or more facing breakends (excluding LINE & assembled & simple non overlapping DEL/DUP) IF the major allele copy number in the segment immediately after the facing breakend is lower than the breakend ploidy, after discounting facing breakends in the original cluster. In the case of there being more than 1 facing breakend, the highest ploidy breakend is clustered and the process repeated. This clustering is limited to a proximity of 5 million bases and bounded by the centromere, since although more distal events are on the same chromosome may be definitely on the same derivative chromosome, this does necessarily imply they occurred concurrently.

*Local Overlap*

Merge any clusters or SVs where each cluster has either an inversion or a DEL exceeding the LONG_DEL_LENGTH, or a DUP exceeding the LONG_DUP_LENGTH and they overlap or enclose each other AND the 2 variants have at least one pair of breakends not more than 5M bases apart that either face each other or form a deletion bridge.

*Foldbacks on same arm*

Merge any 2 clusters with a foldback on the same chromosomal arm.

Resolve DEL, DUP and consistent 2-break junction events

At this step, DEL, DUP and reciprocal clusters are resolved to prevent them from over clustering with later clustering rules. Specifically, the following types of events are resolved:

- **Deletions** - Simple deletions less than the LONG_DEL_LENGTH
- **Tandem duplications** - Simple duplications less than the LONG DUP_LENGTH
- **Reciprocal events and 2-break junction templated insertions -** Pairs of overlapping inversions or translocations where the deletion bridge and/or overlap at both breakends are less than the LONG_DEL_LENGTH / LONG_DUP_LENGTH.

We also allow for synthetic variants of these events to be resolved, where the variants create a DEL, DUP or reciprocal event with the same geometry but with one or more shards inserted. All of these simple and synthetic clusters types are excluded from all subsequent clustering rules.

Common arm clustering rules

*Translocations with common arms*

Merge any 2 unresolved clusters if they touch the same 2 chromosomal arms. SVs which link the 2 arms but are in a short templated insertion (< 1kb) are ignored.

*Single breakends on same arm with matching satellite repeat type clustering*

Where complex events touch satellite repeats we frequently find many single breakends on the same chromosome with links to the same type of repeat. In particular this can occur when shattering events include complex focal scarring in centromeric regions leading to many unresolved single breakends

We therefore merge any cluster with less than or equal to 1 non single breakend and non inferredbreakend with any other cluster which contains a single breakend on the same chromosome with matching repeat class or type for the following cases:

- RepeatClass = 'Satellite/centr' (centromeric)
- RepeatType = '(CATTC)n' (satellite repeat type)
- RepeatType = '(GAATG)n' (satellite repeat type)
- RepatType = 'HSATII' (pericentromeric)

To protect against false positives and joining complex clusters which both touch repeats, but otherwise don't appear to overlap, we avoid clustering 2 clusters which already have multiple non single breakends.

We don't cluster other common sequences such as telomeric sequences, Sine/Alu or LINE/L1 as these tend to be associated with genome wide insertion patterns rather than specific clusters which touch a repetitive region.

##### Incomplete and small complex cluster merging

These rules are implemented to merge small unresolve with 3 or less variants to other unresolved clusters with an arbitrary cluster size where the location and orientation of proximate or overlapping breakends between the 2 clusters indicated that they may be linked.

*Breakends straddled by consecutive same orientation breakends*

Merge any non resolved breakend to a cluster which straddles it immediately on both sides with 2 breakends facing the same direction, and where the facing breakends have matching ploidy.

*Breakends straddled by foldbacks*

Merge any non resolved breakend into a cluster which has 2 different foldbacks straddling it immediately on both sides and at least one of the foldbacks faces the breakend.

*Extended chainable proximity for complex and incomplete events*

Merge any neighbouring non resolved clusters that are within 5M bases and which have facing flanking breakends on each cluster which could form a templated insertion with matching ploidy. In the case of a foldback the ploidy of the facing breakend is also permitted to match 2x the ploidy.

#### 3. Chaining of derivative chromosomes

A chaining algorithm is used to predict the local structure of the derivative chromosome within each cluster. The chaining algorithm examines each cluster independently and considers all possible paths that could be made to connect facing breakends into a set of continuous derivative chromosomes. LINE insertion type clusters are excluded from chaining.

##### Overview of chaining model

A 'chain' in LINX is modelled as a set of 1 or more structural variants connecting a pair of breakends with a ploidy and a ploidy uncertainty. Initially each cluster begins with 1 chain for every SV in the cluster. LINX iteratively makes 'links' between chain ends, resolving 2 of the chains into 1 combined chain and determining a combined ploidy and ploidy uncertainty for the new combined chain from the 2 constituent chains. The order of linking of chains is prioritised, such that the most likely linkages are made first. Structural variant calls from GRIDSS that are shown to be linked empirically by GRIDSS are linked first followed by a set of heuristics to prioritise the remaining uncertain links. This process continues to extend the length of and reduce the number of chains until no further links can be made.

*Constraints on linking breakends*

In general each pair of facing breakends is considered by LINX as a candidate chain. However, 3 key constraints are applied in limiting candidate pairs:

- **'Trans' phased breakends are not permitted to chain -** A breakend may not be chained to another breakend within it's anchoring supporting distance unless it is linked by assembly.

- **Closed loops without centromeres are only allowed for double minutes** - With the exception of double minutes (see below), derivative chromosomes are assumed to connect to either a centromere or telomere at each end. Hence, chains which do not cross a centromere are not allowed to form closed loops (ie. the 2 breakends flanking a chain are not allowed to join to each other) except when specifically identified as a double minute.

- **A chain should not pass through a region with lower available chromatid copy number than the ploidy of the chain** - Chains are generally assumed to take place on a single chromatid except in the rare case that 2 distinct chromatids of the same chromosome are . The copy number for the affected chromatid allele is calculated for each segment adjacent to each cluster by first determining the copy number of the unaffected allele across each contiguous set of breakends in the cluster and subtracting that from the total copy number. The total ploidy of linked chains crossing a segment should not exceed the calculated chromatid copy number for that segment. In some noisy copy number regions or where both chromatids are involved we may not be able to determine an undisrupted allele and the chromatid copy number is not calculated. Additionally, if all possible links have been exhausted using this rule, the rule is relaxed such that chaining can continue under the assumption that allele specific copy number may not have been estimated accurately

*Uniform ploidy clusters*

Chains in LINX are a walkable set of linked breakends with a common ploidy. Uncertainty in ploidy measurement is one of the key confounding factors in predicting the derivative structure. Many clusters however do have the same ploidy for all variants and these uniform ploidy clusters are significantly simpler to chain into a derivative chromosome as each pair of breakends can only be linked with a single ploidy.

Hence, each cluster is tested to see whether all its SVs could be explained by a single ploidy value, using each SV's ploidy estimate and range. Specifically if all SVs have a ploidy range which covers the same value, even if not an integer, then the cluster is considered to be of uniform ploidy and not replication will occur in the chaining routine. Furthermore, if all SVs have a ploidy min less than 1 and a max > 0.5, then the cluster is also considered uniform.

*Variable ploidy clusters*

For all clusters that cannot be resolved by a single uniform ploidy, further considerations apply to explain the amplification of parts of the cluster. Biologically, each SV initially occurs by joining 2 breakends with a ploidy of 1. However, a single chain in a cluster may contain SVs with different ploidies as a result of a replication from either a foldback inversion in a breakage fusion bridge event, or a later tandem duplication of part of the derivative chromosome. In these scenarios, the duplicated variants will appear multiple times in a single chain either repeated in the same direction in the case of a duplication or inverted in the case of a foldback.

Duplication events are permitted to 'replicate' a chain in 2 different ways:

1. **Foldbacks:** Foldbacks with half the ploidy of another chain are permitted to link both their breakends to the same breakend of that chain, making a new chain of half the ploidy with the other unconnected breakend of the non foldback chain at both ends of the new chain. In this foldback replication case the new chain may also be treated as a 'chained foldback' and extended further in the same manner if possible.

2. **Complex Duplications:** Conversely, complex duplications with half the ploidy of another chain are permitted to join both of their breakends simultaneously to either ends of the other chain, effectively duplicating the entire chain, but keeping the same start and end breakends with half the ploidy.

Partial replication of a chain is also possible by later variants which affect chains that have also been duplicated. In this case the higher ploidy chain is split into 2 separate chains, one which is linked to and given the ploidy of the lower ploidy chain, and the other which is given the residual ploidy.

Implementation of chaining algorithm

Linx first resolves all assembled SVs in the cluster into chains. LINX then keeps a cache of a set of chains consisting initially of all 'single variant' chains (ie. lone SVs) and assembled chains. Each chain has 2 breakends, a ploidy and a ploidy uncertainty. Each potentially linkable pairs of facing breakends (subject to the available chromatid copy number and no closed loops rules described above) is also cached as potentially linked chains.

The following steps are then applied iteratively to join chains together until no more chain links can be made:
1. Apply priority rules below to choose the most likely linked pair of chains from the cache
2. Merge chains:
    a. Create new combined chain and calculate the ploidy & ploidy uncertainty
    b. For replication events, replicate and halve the ploidy of the chain, inverting if it is a foldback type event.
    c. In the case of partially split chains, the higher ploidy original chain is kept with it's residual ploidy
    d. Remove merged chains
3. Update cache of linked breakend pairs that include a breakend on the merged chains

*Prioritisation of chain links*

Where more than 1 possible pair of linkable chain exists in the cache, links are ranked and chosen by the following criteria in descending order of importance:

1. Links with available chromatid copy number
2. Links containing highest ploidy foldback or complex duplication chain
    a. Links where it splits another chain with 2x its ploidy
    b. Links where it matches the ploidy of another chain
    c. Links where it splits another foldback with greater than 2x ploidy
    d. Links where it is itself split by another foldback or complex duplication with half the ploidy
3. Breakends with a single link possibility
4. Links with highest matching ploidy status (MATCHED > OVERLAPPING_PLOIDY_RANGE > NO_OVERLAP)
5. Adjacent links

6. Higher ploidy links (allowing for 0.5 abs and 15% threshold)
7. Shortest link

For uniform ploidy clusters, only rules 1, 3,5 & 7 are considered in the prioritisation of links.

A cluster can be chained into more than 1 chain, each one representing the neo-chromosomes resulting from the rearrangement event.

Chaining is often imperfect and incomplete due to the inclusion of single breakends and uncertainty about ploidy and subsequent breakend replication.

*Special considerations for double minutes*

Double minutes and the SVs contained within them subject to special chaining rules in LINX. They are therefore identified prior to chaining taking place. The key principle used to identify double minutes is to look for high ploidy SVs that cannot easily be explained by other processes, particularly breakage fusion bridge.

Specifically the criteria for a double minute are:

- There is at least one breakend with a ploidy of at least 8 AND at least 2.3x the adjacent major allele copy number
- The highest ploidy cannot be explained by a set of foldbacks which form a breakage fusion bridge event. A breakage fusion bridge event is considered plausible if the sum of foldback ploidies + half the maximum inferred breakend ploidy (which could plausibly be a missed foldback) is greater than or equal to the maximum ploidy AND at least one foldback has a ploidy of at least 15% of the maximum ploidy. Even if these criteria are met, if one of the foldbacks is dominant (>75% of the foldback ploidy) then it cannot have been amplified by other foldbacks and breakage fusion bridge is still marked as implausible.

If a cluster is identified as containing a potential double minute then LINX will attempt to first chain the high ploidy variants in that cluster into a closed loop. If a closed loop cannot be made, then chaining proceeds as normal. If a closed loop can be made then any SVs in the cluster with both breakends fully contained within a closed double minute can only link to other variants within the double minute OR to the double minute forming variants. These links are likely lower ploidy disruptions which occurred after the double minute was first replicated are present on a subset of the double minute chromosomes.

##### Chain annotations

The following data is captured for each TI:

- Whether the link is assembled
- Distance to the next link and whether it traverses any other breakends or links
- Overlap with a gene and any exon matches (eg a pseudogenes)

*Annotation of local topology*

Consecutive breakends with no more than 5kb between them or which are part of the same foldback inversion are grouped together into a local topology group and given an id. The number of TIs formed by chained segments wholly within the local topology group are counted and a topology type is given to the remaining variants based on the breakend orientations. The topology types are categorised as one of the following (after excluding all TIs)

- TI_ONLY - All breakends in the group form templated insertions
- ISOLATED_BE - One breakend only

- DSB - A par of breakends forming a deletion bridge
- FOLDBACK - One foldback only
- FOLDBACK_DSB - A foldback with the outer breakend forming a deletion bridge
- SIMPLE_DUP - A single DUP of <5k bases
- COMPLEX_LINE - Any other cluster that is resolved as LINE
- COMPLEX_FOLDBACK - Any other cluster that includes a foldback
- COMPLEX_OTHER - Any other cluster

#### 4. Gene impact & fusion prediction

Annotation of breakends with potential gene impact

For each breakend we search for genes that could be potentially disrupted or fused by the structural variant.

To do this we find and annotate the breakend for any transcript that either:
- Has an exon or intron overlapping the breakend
- Has it's 5' end downstream of and facing the breakend and less than 100k bases where no other splice acceptor exists closer to the breakend.

Each breakend is additionally annotated for the transcript with the following information:
- **disruptive**: disruptive for a particular transcript if the SV it belongs to is an inversion,single breakend or translocation or if the SV overlaps at least one exon in the transcript AND the variant is not part of a chain which does not disrupt the exon ordering in the transcript AND the variant is not resolved LINE insertion
- **transcript coding context**: UPSTREAM, 5_UTR, CODING, 3_UTR, DOWNSTREAM OR NON_CODING
- **Gene orientation**: relative orientation of gene compared to breakend (UPSTREAM or DOWNSTREAM)
- **exonic** (TRUE/FALSE)
- **exact base phase**: The exact base phasing of the current location
- **Next splice site information**: The distance to, phasing of and exon rank of the 1st base of the next facing splice acceptor or donor (note: phasing will be different from exactBasePhase if the breakend is exonic in the transcript or the coding context is upstream. Null if there are no subsequent splice sites in the gene.
- **exon total count**: The total number of exons in the transcript (for reference)
- **transcript biotype**: The ensembl biotype of the transcript

Fusions are predicted by looking for consecutive and novel splice donor-acceptor pairings that are joined together in derivative chromosomes by either a single structural variant or a continuous chain of structural variants.

Fusion prediction

*Curate a list of known fusions*

A list of 393 curated known fusion pairs was sourced by taking the union of known fusions from the following external databases:
- Cosmic curated fusions[12] (v83)
- OncoKb[9] (download = 27-Nov-2018)
- CGI[8] (download = 27-Nov-2018)
- CIViC[7] (download = 27-Nov-2018)

We then also created a list of promiscuous fusion partners using the following rules

- **3’ promiscuous:** Any gene which appears on the 3’ side in more than 3 of the curated fusion pairs OR appears at least once on the 3’ side and is marked as promiscuous in either OncoKb, CGI or CIVIC
- **5’ promiscuous:** Any gene which appears on the 5’ side in more than 3 of the curated fusion pairs OR appears at least once on the 5’ side and is marked as promiscuous in either OncoKb, CGI or CIVIC

*Identify viable fusions*

For each single SV and for every facing pair of SVs in the same chain identify all viable fusions which satisfy the following conditions:

- Chain must join appropriate contexts of the 5’ and 3’ genes after allowing for any skipped exons (see table below)
- Chain must be disruptive to the 5’ partner
- Not disrupted by an intermediate splice acceptor / splice donor. Exceptions:
    - Traversal can be valid even if there are intermediate splice acceptors as long as upstream partner, downstream partner and all intermediaries are 5’ non-coding)
    - Splice donors/acceptors can be skipped within the same gene if there is no in-frame valid fusion without skipping (this is to deal with alternative splicing). LINX reports the number of exons skipped on both the 5’ and 3’ transcript.
- Not terminated on the 5’ partner side by a chained breakend prior to the start of the 5’ gene
- Not terminated on the 3’ partner side by a chained breakend prior to the last coding base of the 3’ gene.
- For chained fusions, the total chain length must be less than 100kb
- For upstream context, 10kb up from specific transcript start or 100kb for known fusion pairs

The following table shows allowed contexts:

<table>
<tr><td></td><td colspan="6">3’ Fusion Partner Context</td></tr>
<tr><td>5’ Partner Context</td><td>Upstream</td><td>5’UTR Intronic</td><td>5’UTR Exonic</td><td>Coding Intronic</td><td>Coding Exonic</td><td>3’ UTR or Non-Coding</td></tr>
<tr><td>Upstream</td><td>X<br>(Enhancer)</td><td colspan="4">X<br>(promoter loss)</td><td rowspan="4">X<br>(No Downstream Impact)</td></tr>
<tr><td>5’UTR or non-coding Intronic</td><td>YES(1)</td><td>YES</td><td>YES</td><td colspan="2">X<br>(4)</td></tr>
<tr><td>5’UTR or non coding Exonic</td><td colspan="2">X<br>(3)</td><td>YES</td><td colspan="2">X<br>(4)</td></tr>
<tr><td>Coding Intronic</td><td>YES<br>(1)</td><td colspan="2">SEE NOTE<br>(5)</td><td>YES</td><td>YES<br>(2)</td></tr>
</table>

| | | |
|---|---|---|
| Coding Exonic | X<br>(3) | YES |
| 3'UTR | X<br>(post stop codon in upstream gene) | |

(1) for breakends in the upstream region of the 3' partner, the 2nd exon (1st splice acceptor) is checked for phasing and assumed to be fused exon. 5' partner coding to 3' partner upstream is also possible if the 3' partner coding region starts in the 1st exon and the 1st splice acceptor in the 2nd exon is in phase
(2) If fusing intron to exon, will fuse with the next downstream exon, so check against frame of end of exon instead of exact base.
(3) Exonic to Intronic can occur if alternative splicing causes exon with exonic breakend to be skipped
(4) 5' partner 5'UTR or non-coding to coding region of 3' partner could technically make a fusion, but would need to find 1st alternative start codon also in-frame.
(5) Coding Intronic to non-coding allowed only when transcript starts on 1st base of next downstream exon - in this case we fuse to the first base of the gene which is allowed.

*Prioritise transcripts*

Each candidate fusion gene pair may have many potential fusion transcripts on both the 5' gene and 3' gene.

LINX will prioritise transcripts and choose one fusion per cluster per fusion gene pair by the following rules in order:

1. Viable non-disrupted fusion
2. Transcript with In-frame fusion without skipped exons
3. Both the 5' and 3' transcripts are canonical
4. Best 3' partner, ranked by canonical and then longest (non NMD) protein coding
5. Best 5' partner out of those ranked by canonical, then longest protein coding

*Determine reportability*

Mark as reportable based on the following rules

1. Matches an exact fusion from the curated list OR (is intergenic AND (matches 5' promiscuous OR matches 3' promiscuous gene) AND no exons are skipped)
2. Curated domains which must be preserved
    a. Ets domain
    b. Protein kinase domain
    c. Epidermal growth factor-like domain
    d. Ankyrin repeat-containing domain
    e. Basic-leucine zipper domain
    f. High mobility group box domain
3. Curated domains which must be disrupted if exist
4. Raf-like Ras-binding (mainly affects BRAF)
5. For intragenic fusions, must start and end in coding regions of the gene
6. 3' partner must be a protein coding gene and the transcript does not result in nonsense mediated decay

Amplification, deletion and disruption drivers

*Homozygous disruption drivers*

LINX can optionally take as input a catalog of point mutation, amplification and homozygous deletion drivers which is created by PURPLE based on the raw somatic variant data and determined copy number profile. LINX leverages it's chaining logic to extend the driver catalog by searching for 2 additional types of biallelic disruptions which disrupt all copies of the gene but do not cause a homozygous deletion in an exonic segment (which is

PURPLE’s criteria for homozygous deletion). Specifically LINX searches for 2 additional types of homozygous disruptions:

- **Disruptive breakends -** Any disruptive breakend which causes the intronic copy number to drop to <0.5 copy number after allowing for the ploidy of any overlapping deletion bridges.
- **Disruptive duplications -** Any duplication which has both breakends disruptive in the transcript and a ploidy >= flanking copy number at both ends end.

*Linkage of drivers to contributing structural variant clusters*

We link each driver in the catalog that is affected by genomic rearrangements (ie. high level amplifications, homozygous deletions and biallelic point mutations in TSG with LOH and the homozygous disruptions found by LINX) to each structural variant cluster which contributed to the driver. 1 or more structural variant clusters may contribute to each event and/or the driver may be caused by a whole chromosome or whole arm event which cannot be mapped to a specific variant but which has caused significant copy number gain or loss

Amplifications drivers are linked to either one or more clusters which explain the gain in copy number over the gene (marked as type 'GAIN'), a gain in copy number in the centromere (marked as type 'GAIN_ARM') or across the whole chromosome (marked as type 'GAIN_CHR'). More than 1 of these factors can be recorded as a cause of amplification if its copy number gain is at least 33% of the largest contributing factor. To determine whether a cluster contributes to gene amplification, the copy number change of all its breakends surrounding the gene are summed into a net cluster copy number gain, and the copy number loss of any opposing clusters are subtracted. If a net gain remains, then the cluster is considered as contributing to the gene amplification.

Homozygous deletions drivers are linked to clusters that either directly bound by the homozygous deleted region (svDriverType = DEL) or a cluster that bounds the LOH (svDriverType = LOH). Biallelic point mutations drivers in TSG with LOH are linked only to the cluster that bounds the LOH (svDriverType = LOH). If the LOH bounds extend to the whole arm or whole chromosome, then a record is created with linked clusterId is set to NULL and the svDrivertype is marked as LOH_ARM or LOH_CHR respectively.

The supported linkages between drivers and SVs are summarised in the table below

| Driver Type | Events per driver | svDriverTypes |
|---|---|---|
| Amplification | 1+ | GAIN<br>GAIN_ARM<br>GAIN_CHR |
| Homozygous Deletion | 2 | DEL<br>LOH<br>LOH_ARM<br>LOH_CHR |
| Biallelic point mutation in TSG | 1 | LOH<br>LOH_ARM<br>LOH_CHR |

#### Visualisation

LINX provides functionality to present detailed visualisation of genomic rearrangements including genic impact in CIRCOS [28] format. LINX writes a set of 'VIS' files in a specific format which form the base data to generate the visualisation. The visualisation tool only depends on these files and so in principle any tool could provide SV & CN data in this format.

There are 3 main components in the output of the LINX visualisation each described below.

##### CIRCOS view

The CIRCOS view shows either the complete set of rearrangements in a cluster or set of clusters if 1 or more cluster ids are specified ('cluster mode') or all clusters that touch a chromosome if a chromosome is specified ('chromosome mode').

The CIRCOS view has 6 tracks showing from innermost to outermost (figure 4a):

1. Break junctions
2. Minor allele copy number profile
3. Copy number profile
4. Derivative chromosomes
5. Impacted genes (if any)
6. Affected chromosomes

The scaling of both distances and ploidies in the figure has been modified to make the figure readable. Specifically, the distances between each feature in the chart (either breakend or gene exon start or end) are modified to a log based scale, so that the entire genomic rearrangement spanning millions of bases and multiple chromosomes can be viewed, but that local topology of regions with high densities of breakpoints can be introspected. Ploidy is set with a linear scale but is scaled down if the maximum cluster ploidy exceeds 6 or if the total density of events exceeds a certain maximum to ensure that even the most complex clusters can be introspected. Additionally, if the total number of breakends displayed exceeds XX then the break junctions become increasingly transparent such that other features on the plot don't become obscured.

Another key feature of the CIRCOS plot is the ability to trace the derivative chromosome(s). Each segment in the 4th track represents a segment of the derivative chromosome and is linked on both ends either to a centromeric or telomeric end (marked with an open or closed square respective) or a breakend (marked with a track or triangle in the case of foldbacks). Each derivative chromosome can be traced continuously from one telomeric / centromeric end to another (or to a single breakend if one is reached) by following a continuous series of segments and breakends. To make this easier to follow, each time a new segment is connected on a chromosome the segment is offset outwards slightly. Hence the derivative chromosomes can be traced from the inside to the outside of the 4th track of the diagram. A cluster may contain 1 or more derivative chromosomes. In cluster mode each derivative chromosome will be shown in a different colour for ease of viewing (with a maximum of 10 colours after which all derivative chromosomes are shown in black). In chromosome mode, derivative chromosomes will be shown in the same colours, but each cluster is shown in a different colour. Red and green are reserved for simple deletions and tandem duplications respectively. Since there may be many of these on a single chromosome, telomeric and centromeric connectors are not shown for these simple variant types. Light blue is also reserved for LINE clusters which can also be frequent in samples with highly deregulated LINE machinery.

Most of the possible annotations are shown in the LINX visualisation guide [see Figure 4a]. Additionally, 2 types of genomic regions which are frequently disrupted in tumor genomes are indicated using light grey shading on the copy number regions in the 3rd track. For known LINE source elements the green copy number section is shaded and for known fragile sites the red copy number section is shaded light grey.

#### Chromosome view

Since the CIRCOS only represents a part of the genome, the chromosome view is provided to indicate which parts of the genome is shown. Each of the chromosomes included in the cluster(s) shown is displayed. The part of the chromosome that is included in the figure is highlighted in the colour matching the colour used in the outer ring of the CIRCOS. The banding and location of the centromere on each chromosome is also shown.

An example of the chromosome view is shown below indicating the cluster includes a large section of chromosome 7 including the centromere and a small slither of chromosome 15 on the Q arm:

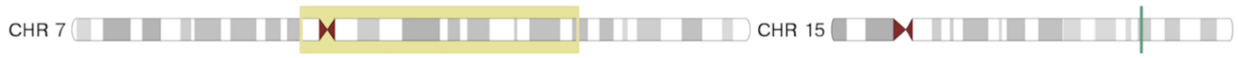

#### Fusion view

The fusion view is added for reportable fusions only in LINX. It's purpose is to show the predicted structure of the fused gene. The fusion includes the fused segments of both the 5' and 3' partner in blue and red and always reads from left to right. The gene representation for each genes and follows the standard conventions of thick bands for coding regions, thinner bands for 5' UTR and 3' UTR exonic regions and thin lines to represent the intronic sections. Protein domains are shown in coloured bands across the exons which they include and are labeled in the accompanying legend. As with the CIRCOS view the lengths of exonic gene segments in the fusion view are scaled by a log scale to improve readability. The intronic segments are set to a fixed segment length regardless of the length.

The fused gene is shown up to and including the breakend on either side that is connected either directly in the case of a simple fusion or via a chain in a chained fusion. If LINX predicts that one or more exons are skipped, then the skipped exonic segments and protein domain sections are faded.

For example in the following TMPRSS2-ERG fusion, exons 3, 4 & 5 are faded and the LDL domain is also faded, indicating that LINX predicts these exons are skipped in order to make a viable in-frame protein, despite the break end occuring after the 5th exon:

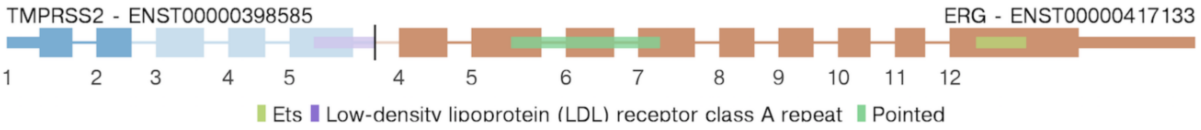

A similar case can be seen in this example where the 5'UTR region of NDRG1 is fused upstream of the 1st exon in PLAG1. Since the 1st exon of a gene has no splice acceptor, LINX predicts the 1st exon is skipped and it is faded on the chart with the fusion connecting to the start of exon 2 which also begins in the 5'UTR region of PLAG1:

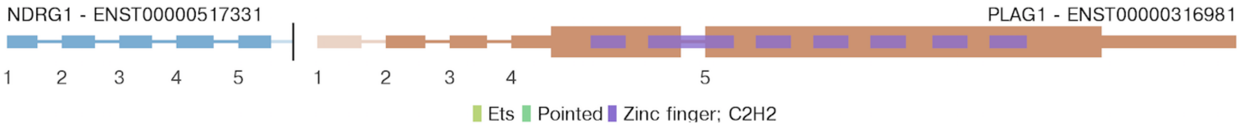

Even when no exons are skipped, a protein domain may be partially disrupted if it extends to an exon the other side of the breakpoint. In the below example, you can see the SH2 protein domain extends before exon 16 and is faded and disrupted:

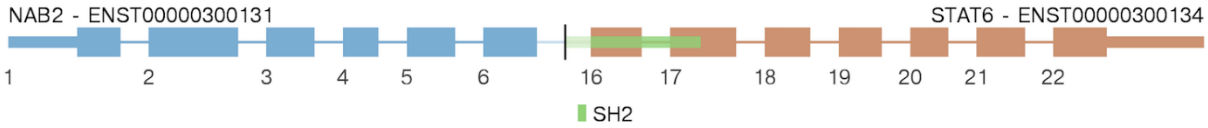

Whilst fusions are normally intronic, rare exonic to exonic fusions do occur. The below figure shows a CIC-FOX04 example where the 2 exons are directly fused

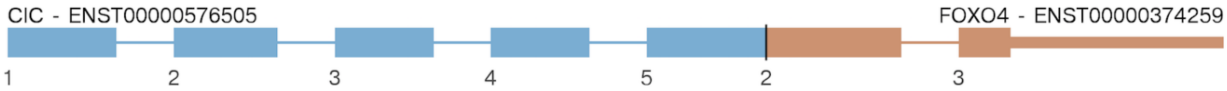

### Validations

#### Benchmarking on COLO829T and NA12878T

##### GRIDSS structural variant comparison on NULL tumors & COLO829T

The NA12878 and COLO829T/COLO829BL cell lines (ATCC® CRL-1974™ and 1980™ respectively) were each sequenced three times to 100x/40x using the HMF workflow [19] and aligned against hg19 without alt contigs using BWA 0.7.15. Manta 1.5.0 was run with default parameters and GRIDSS run as outlined above.

The COLO829 somatic SV truthset was generated using an orthogonal sequencing strategy. We sequenced the COLO829BL and COLO829T cell lines using Illumina HiSeqX (ENA run accessions ERR2752449 and ERR2752450 for COLO829BL and COLO829T, respectively), Oxford Nanopore (ERR2752451 and ERR2752452), Pacific Biosciences (ERR2752447 and ERR2752448) and 10X genomics (ERR2820166 and ERR2820167). All data is grouped under ENA study accession PRJEB27698.
Raw data was analysed for structural variants using the following tools:

- Illumina data was mapped using BWA 0.7.5 and SV calling was performed with GRIDSS 2.0.1 and somatic SVs were filtered using gridss_somatic_filter.R.
- Nanopore data was mapped using NGMLR 0.2.6 and SV calling was performed with both NanoSV 1.2.0 and Sniffles 1.0.8 separately for COLO829T and COLO829BL. All SVs were merged with an overlap window of 200 base pairs using SURVIVOR 1.0.6 and SVs not present in COLO829BL were kept.
- Pacbio data was mapped using minimap2 2.11-r797 and SVs were called using pbsv 2.1.0 in joint calling mode for COLO829T and COLO829BL. Only SVs with no evidence in COLO829BL were kept.
- 10X genomics data was processed using Longranger 2.2.2 with default settings for COLO829BL and somatic mode for COLO829T. SV calls for both cell lines were merged with an overlap window of 200 base pairs using SURVIVOR 1.0.6 and SVs not present in COLO829BL were kept.

Somatic SV calls for each technology were merged with an overlap window of 200 base pairs using SURVIVOR 1.0.6. and all candidate breakpoints were subjected to independent validation by targeted capture and/or PCR-based approaches. SVs detected with two or more techniques that failed in these validation experiments were curated by manual inspection of the mapped reads using IGV (ref). A total of 69 SVs were finally considered as true somatic SVs for COLO829T.

Call matching was performed using the StructuralVariantAnnotation BioConductor package (DOI 10.18129/B9.bioc.StructuralVariantAnnotation) with a 4bp margin on top of any caller-reported microhomology allowed when considering calls to match. Single breakends calls were considered true positives if the position and orientation matched any truth set breakpoint and for the purpose of calculating sensitivity, were considered half a match to the truth breakpoint.

##### Comparison of PURPLE to other copy number tools

To compare the performance PURPLE to other CNV/SV tools, CONSERTING, WEAVER, and ASCAT were run on COLO829 tumor/normal cell line pair using a no alt hg19 reference genome. Weaver 0.21 was run according to the instructions in README.rst with with a tile size of 1000. CONSERTING was run as per the provided README_2016_v2.pdf documentation. A window size of 100 was used and the recommended but optional LOH and WGS CREST steps were run. Average copy number reported by ASCAT was used to convert CONSERTING mean coverage to copy number estimates.
Copy number segment bounds were matched to the nearest SV with 100,000bp from the corresponding SV call set (PURPLE/GRIDSS, CONSERTING/CREST, Weaver/Weaver, ASCAT/GRIDSS) regardless of breakpoint orientation. Copy number consistency across breakpoints was determined by the absolute difference between the copy number delta across each side of the breakpoint, normalised by the maximum copy number of the segments matched to the breakpoint. Breakpoint and copy number transitions that did not have an exact position matches after taking into account any breakpoint micro-homology were excluded.

#### Validations on Hartwig Medical Cohort

##### Cohort level results

All cohort level validations and results in this paper have been produced from running the GRIDSS, PURPLE and LINX toolkit on a pan-cancer cohort of 3,524 unique patient samples from the Hartwig Medical Foundation cohort which has been previously described [19]. This includes the analysis CIS vs TRANS breakends, the prevelance of single breakend structural variant calls and the concordance of copy number and structural variants. 534 of the 3,524 samples have matching whole transcriptome RNA data which we have use for fusion validation as described below.

##### RNA validation of LINX fusion predictions on 534 samples

STAR-Fusion (v1.5) was run for 534 samples with matching whole transcriptome data to validate the SV fusion predictions, producing a total of 1,478 predicted in-frame fusion pairs. Out-of-frame RNA fusions and fusions to non-coding downstream partners or the 3' UTR regions in the downstream partner were ignored as they do not make a viable protein. Fusions in the 3'UTR region of the upstream partner were also ignored. We also filtered for recurrent artefacts across many samples, and deduplicated gene pairs found multiple times to choose one fusion prediction per gene pair per sample.

The in-frame LINX fusion predictions where matched to the RNA predictions if the same gene pair was predicted and the locations of the LINX breakends were consistent with the fused RNA locations (ie. the breakend must be upstream of the RNA fused location on the 3' gene and downstream of it on the 5' gene). Intragenic LINX fusions (ie exon deletions or duplications) were excluded from the analysis as they are not called by STAR-Fusion.

Altogether 482 of the 534 samples had at least one inframe fusion prediction from either RNA or DNA (supplementary table 3).

#### Capture panel validation structural variant calling on 13 tumor samples

13 of the 3,524 samples were selected for capture panel validation of the structural variant calls. Each variant called in GRIDSS was compared with variants called from Manta (for variants longer than 50 bases) and/or Strelka (for variants from 32-50bases in length) to determine if the variant is shared or private according. Variants were marked as matching GRIDSS if the start and end chromosomes and orientation both match and start end positions (including confidence intervals) were within 20 bases of each other.

A probe was created for each of the shared and private variants. For each (non-SGL) variant we created 3 probes of 120 bases each. Two reference probes leading up to the breakends as well as another SV probe going through the structural variant. The reference probes were designed to end 20 bases before the start of each structural variant breakend. The SV probe consists of any insert sequence flanked by equal amount of bases from each side of the structural variant. For each SGL variant we created 2 probes. One reference probe as described above and one SV probe which includes no more than 60 bases of the insert sequence and the remainder coming from the reference leading to the break point.

Together, this created a total of 17,125 probes of 120 nt in length targeting 5,821 break-junctions (see supplementary table 1) which were ordered as custom target capture probes at Twist Biosciences (catalog ID 100533). For each of the 13 samples 50 ng input DNA was used for indexed library construction with enzymatic fragmentation (Twist kit catalog IDs 100253, 100255 and 100401) according to the manufacturer's protocol. With a bead-based size selection after PCR to remove the remaining larger fragments (>700bp). Multiplexed hybridization was performed using the Twist Hybridization (ID100254), Blockers (PN100856) and Wash Kits (PN100214, 100215, 100216)) using Dynabeads™ MyOne™ Streptavidin T1 (Invitrogen PN65604D) following standard Twist protocol. Enriched library molecules were amplified by PCR for 11 cycles and sequenced on the Illumina NextSeq500 2x 150bp High Output run according to manufacturer's standard protocol.

We created a set of predicted alternate contigs from the shared and private structural variant calls using the same technique from above for generating the (non-reference) SV probes and added these to the ref genome. We then mapped each of the reads from the capture panel output with BWA to a hybrid genome including both the GRCH37 reference genome and the novel alternate contig.

We assessed the viability of the probes by mapping each 120 base SV probes to the 2,000 base alternate contigs to determine its mapping quality. Of the 5,821 SV probes, 80 had 0 mapping quality and 5,377 has a perfect mapping quality of 60. Probes with mapping quality less than 20 were ignored as well as 77 micro-satellite probes where we unable to unambiguously validate the length of the expansion with this validation approach. We examined the resultant BAM files for evidence of the SV alternate contigs in the SV source sample BAM as well as the BAM files of each of the other samples as controls for systemic effects for each of the predicted variants. Specifically, the read depth on the alternate contig at the variant location was used to assess the validation status of the variant. Specifically SV calls were marked as validated if all the following criteria are met (and not validated otherwise):

- At least 2 reads were mapped to the alternate contig in the predicted sample
- The support for the alternate contig was significantly higher rate (Poisson model, p=0.001) in the predicted sample than the maximum of the other 13 samples
- <40 reads in total across all 13 control samples were mapped to the predicted alternate contig

#### Third party bioinformatics tools

The following bioinformatics tools have been used directly:

| **Software** | **Version** | **Link** |
|---|---|---|
| bcl2fastq | 2.17-2.20 | http://sapac.support.illumina.com/downloads/bcl2fastq-conversion-software-v2-20.html |
| bwa mem | 0.7.5a- | https://github.com/lh3/bwa |
| sambamba | 0.6.5-0.7.0 | https://github.com/biod/sambamba/ |
| Picard | 1.141-2.18 | https://broadinstitute.github.io/picard/ |
| GATK | 3.4.46 | https://software.broadinstitute.org/gatk/ |
| Strelka | 1.0.14 | https://github.com/Illumina/strelka |
| Manta | 1.0.3 | https://github.com/Illumina/manta |
| STAR-fusion | 1.5 | https://github.com/STAR-Fusion/STAR-Fusion/ |
| ASCAT | 2.52 | https://github.com/Crick-CancerGenomics/ascat |
| dNdScv | 0.1.0 | https://github.com/im3sanger/dndscv/ |
| Circos | 0.69.6 | http://circos.ca/ |
| samtools | 1.2 | https://github.com/samtools/samtools/ |
| snpeff | 4.3s | https://sourceforge.net/projects/snpeff/files/snpEff_v4_3s_core.zip/download |
| vcftools | 0.1.14 | https://vcftools.github.io/index.html |
| bcftools | 1.9 | https://github.com/samtools/bcftools/ |
| SSW Library | 1.0 | https://github.com/mengyao/Complete-Striped-Smith-Waterman-Library |
| mutationalPatterns | 1.4.3 | https://bioc.ism.ac.jp/packages/3.6/bioc/html/MutationalPatterns.html |
| CopyNumber | 1.24.0 | http://bioconductor.org/packages/release/bioc/html/copynumber.html |
| VariantAnnotation | 1.29 | https://bioconductor.org/packages/release/bioc/html/VariantAnnotation.html |
| StructuralVariantAnnotation | 1.0 | https://bioconductor.org/packages/release/bioc/html/StructuralVariantAnnotation.html |
| Gviz | 1.28 | https://bioconductor.org/packages/release/bioc/html/Gviz.html |
| BSgenome.Hsapiens.UCSC.hg19 | 1.4.0 | https://bioconductor.org/packages/release/data/annotation/html/BSgenome.Hsapiens.UCSC.hg19.html |
